# Variation and inheritance of the *Xanthomonas* gene cluster required for activation of XA21-mediated immunity

**DOI:** 10.1101/149930

**Authors:** Furong Liu, Megan McDonald, Benjamin Schwessinger, Anna Joe, Rory Pruitt, Teresa Erickson, Xiuxiang Zhao, Valley Stewart, Pamela C. Ronald

**Affiliations:** Department of Plant Pathology and the Genome Center, University of California, Davis; Research School of Biology, Australian National University, Canberra; Department of Microbiology & Molecular Genetics, University of California, Davis

**Keywords:** *Xanthomonas*, tyrosine sulfating, plant innate immunity

## Abstract

The rice XA21-mediated immune response is activated upon recognition of the RaxX peptide produced by the bacterium *Xanthomonas oryzae* pv. *oryzae* (*Xoo*). The 60 residue RaxX precursor is posttranslationally modified to form a sulfated tyrosine peptide that shares sequence and functional similarity with the plant sulfated tyrosine (PSY) peptide hormones. The five kb *raxX-raxSTAB* gene cluster of *Xoo* encodes RaxX, the RaxST tyrosylprotein sulfotransferase, and the RaxA and RaxB components of a predicted type one secretion system. The identified the complete *raxX-raxSTAB* gene cluster is present only in *Xanthomonas* spp., in five distinct lineages in addition to *X. oryzae*. The phylogenetic distribution of the *raxX-raxSTAB* gene cluster is consistent with the occurrence of multiple lateral transfer events during *Xanthomonas* speciation. RaxX variants representing each of the five lineages, and three *Xoo* RaxX variants, fail to activate the XA21-mediated immune response yet retain peptide hormone activity. These RaxX variants contain a restricted set of missense mutations, consistent with the hypothesis that selection acts to maintain peptide hormone-like function. These observations are also consistent with the hypothesis that the XA21 receptor evolved specifically to recognize *Xoo* RaxX.

## INTRODUCTION

Host receptors activate innate immunity pathways upon pathogen recognition (Ronald & Beutler, 2010). The gene encoding the rice XA21 receptor kinase (Song *et al.*, 1995) confers resistance against most strains of the gamma-proteobacterium *Xanthomonas oryzae* pv. *oryzae* (*Xoo*) (Wang *et al.*, 1996). This well-studied XA21-*Xoo* interaction provides a basis from which to understand molecular and evolutionary mechanisms of host-microbe interactions.

Four *Xoo* genes that are <u>**r**</u>equired for <u>**a**</u>ctivation of <u>**X**</u>A21-mediated immunity, are located in the *raxX-raxSTAB* gene cluster (**Fig. 1**). The 60-residue RaxX predicted precursor protein undergoes sulfation by the RaxST tyrosylprotein sulfotransferase at residue Tyr-41 (Pruitt *et al.*, 2015). We hypothesize that the RaxB proteolytic maturation and ATP-dependent peptide secretion complex (da Silva *et al.*, 2004) further processes the sulfated RaxX precursor by removing its double-glycine leader peptide prior to secretion (Holland *et al.*, 2016). Located outside the *raxX-raxSTAB* gene cluster, the *raxC* gene, an ortholog of the *tolC* gene, encodes the predicted outer membrane channel for this secretion complex (da Silva et al., 2004). Finally, the *raxPQ* genes encode enzymes to assimilate sulfate into 3’-<u>**p**</u>hospho<u>**a**</u>denosine 5’-<u>**p**</u>hospho<u>**s**</u>ulfate (PAPS) (Shen *et al.*, 2002), the sulfodonor for the RaxST sulfotransferase (Han *et al.*, 2012).

**Fig. 1.**
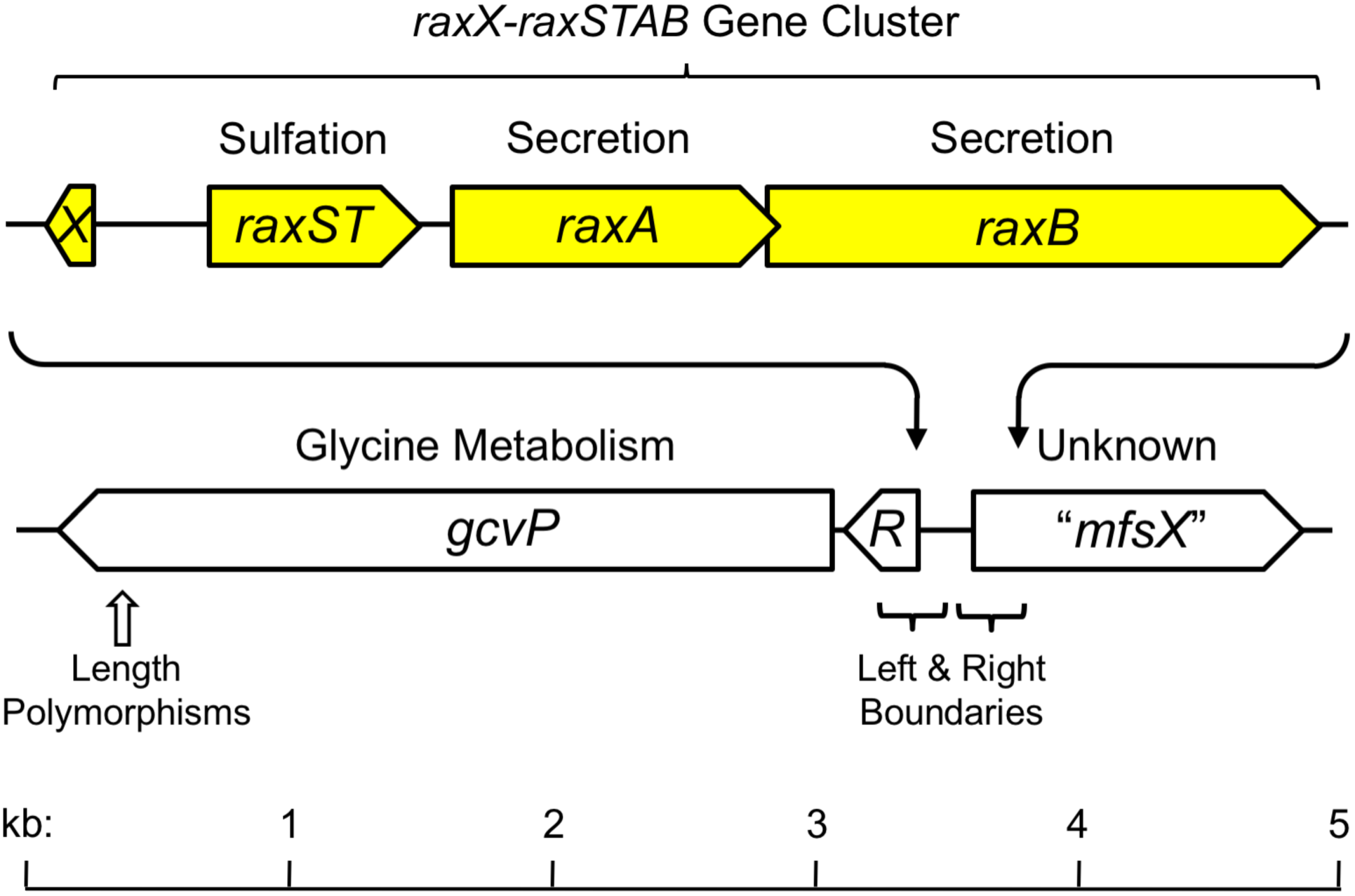
The *raxX-raxSTAB* gene cluster. The *raxX-raxSTAB* gene cluster is located between the flanking *gcvRP* and *"mfsX"* genes. Gene cluster acquisition through lateral transfer is hypothesized to occur by general recombination in the flanking *gcvR* and "*mfsX*" sequences as described in the text. Sequences at the left and right boundaries are shown in **Fig. S2**. Sequences for length polymorphisms in the *gcv*P gene are shown in **Fig. S3**.

In both plants and animals, the post-translational modification catalyzed by tyrosylprotein sulfotransferase is restricted to a subset of cell surface and secreted proteins that influence a variety of eukaryotic physiological processes (Matsubayashi, 2014, Stone *et al.*, 2009). For example, tyrosine sulfation of the chemokine receptors CCR5 and CXCR4 is essential for their functions including as coreceptors for the human immunodeficiency virus gp120 envelope glycoprotein (Farzan *et al.*, 1999, Kleist *et al.*, 2016). In plants, sulfated tyrosine peptides influence cellular proliferation and expansion in root growth, and/or plant immune signaling (Matsubayashi, 2014, Tang *et al.*, 2017). In contrast to these and other examples of protein tyrosine sulfation in animals and plants, RaxX sulfation by the RaxST enzyme is the only example of tyrosine sulfation documented in bacteria (Pruitt et al., 2015, Han et al., 2012).

Mature RaxX is predicted to comprise the carboxyl-terminal residues 40-60, numbered according to the precursor protein (Pruitt et al., 2015, Pruitt *et al.*, 2017). RaxX residues 40-52 share sequence similarity with mature <u>**p**</u>lant peptide containing <u>**s**</u>ulfated t<u>**y**</u>rosine (PSY) hormones (Pruitt et al., 2015, Amano *et al.*, 2007, Pruitt et al., 2017). RaxX, like PSY1, can enhance root growth in diverse plant species (Pruitt et al., 2017). The XA21-mediated response in rice requires residues 40-55 (RaxX16 peptide), whereas plant growth stimulation requires only residues 40-52 (RaxX13 peptide) (Pruitt et al., 2015). **Fig. 2** shows sequences for the RaxX variants examined in this study, together with two representative PSY sequences for comparison.

**Fig. 2.**
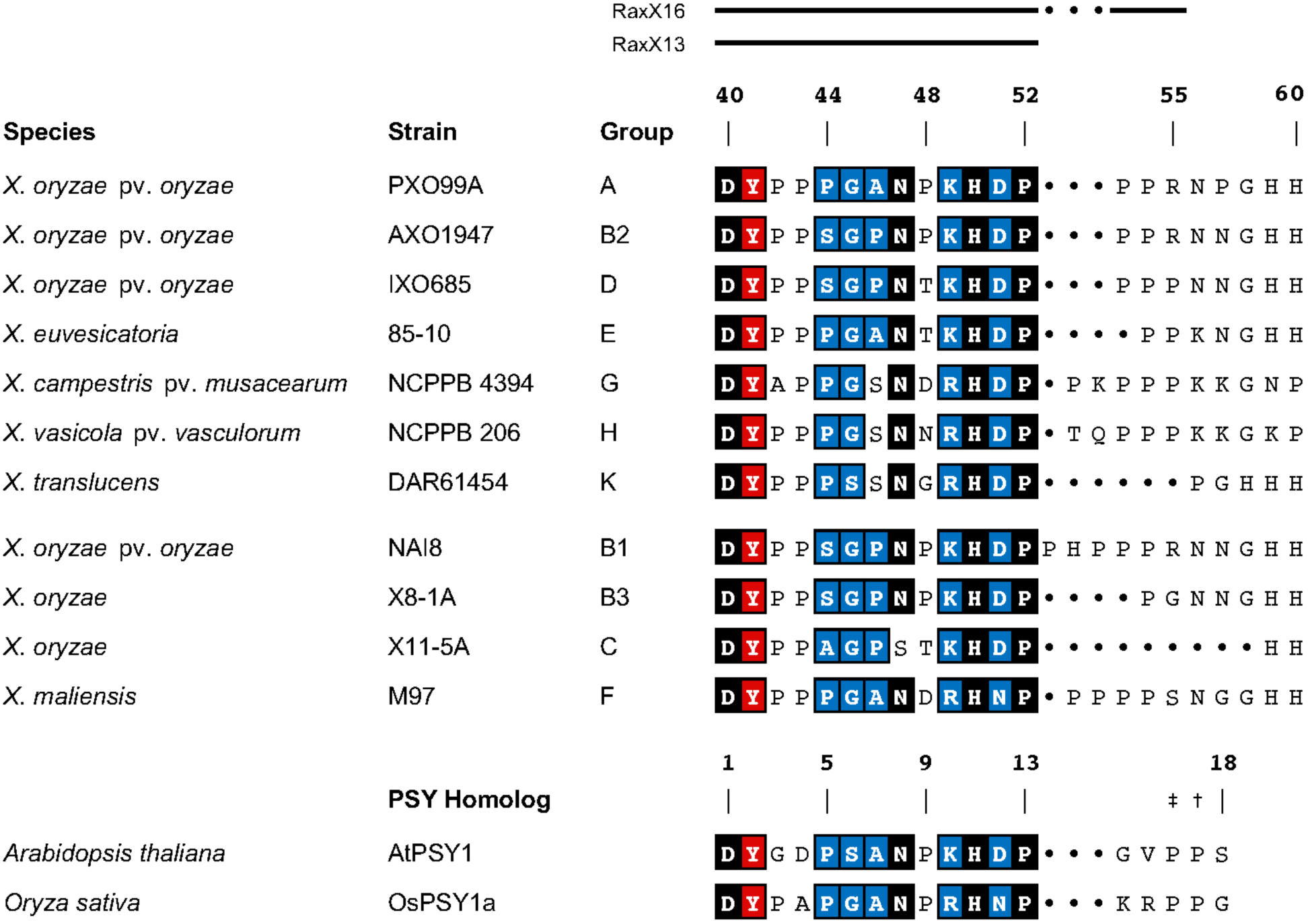
RaxX variants. Sequences show the presumed leader-cleaved forms of RaxX, numbered from the beginning of the precursor sequence. The extent of sequence comprising the RaxX16 and RaxX13 synthetic peptides is indicated above the alignment. Residues are shaded according to conservation in PSY sequences (Pruitt et al., 2017): positions with nearly invariant residues are shaded black, and those with only two or three substitutions are shaded blue. The sulfated Tyr residue is shaded red. Gaps are indicated by dots. Sequence groups are described elsewhere in detail (Pruitt et al., 2017). The subgroups B1-B3 differ only in the carboxyl-terminal sequence beginning with residue 53. *X. oryzae* strains X8-1A and X11-5A are nonpathogenic and therefore do not have pathovar designations. The mature form of *Arabidopsis thaliana* PSY1 (Amano et al., 2007) and the corresponding region from *Oryza sativa* PSY1a (Amano et al., 2007, Pruitt et al., 2017) are shown for comparison. Residues Pro-16 and Pro-17 in AtPSY1 both are hydroxylated [†,‡], and Pro-16 is glycosylated with L-Ara_3_ [‡] (Amano et al., 2007).

RaxX sequences generally are well conserved within different *Xanthomonas* species (Pruitt et al., 2017). In *Xoo* however, RaxX from a strain IXO685, which evades XA21-mediated immunity differs from active RaxX at the critical positions Pro-44 and Pro-48 (**Fig. 2**) (Pruitt et al., 2015). Nevertheless, this RaxX protein stimulates root growth, as do two other RaxX Pro-48 variants from other *Xanthomonas* spp. (Fig. 6 in (Pruitt et al., 2017)).

These results suggest that RaxX recognition by XA21 is restrained by different sequence and length requirements compared to its recognition by the root growth promoting receptor(s) for PSY hormone(s). It also suggests that recognition of RaxX by XA21 is specific to *Xoo*, whereas PSY mimicry is a general feature of RaxX from other *Xanthomonas* spp. Accordingly, we hypothesized that PSY hormone mimicry is the original function of RaxX, whereas immune recognition by XA21 evolved later in response to *Xoo* (Pruitt et al., 2017).

Two general predictions derive from this hypothesis. The first prediction, that PSY hormone mimicry is broadly selective, is supported here by the presence of the *raxX-raxSTAB* gene cluster *Xanthomonas* spp., and by the ability of all RaxX variants tested to stimulate root growth in an assay for PSY function. The second prediction, that recognition by XA21 is restricted to *X. oryzae* lineages, is validated here by the observation that XA21-mediated immunity is not activated by RaxX variants from other *Xanthomonas* spp. These results illustrate how a pathogen protein has evolved to retain its ability to modulate host physiology without being recognized by the host immune system.

## RESULTS

### The *raxX-raxSTAB* gene cluster is present in a subset of *Xanthomonas* spp

We searched databases at the National Center for Biotechnology Information to identify bacterial genomes with the *raxX-raxSTAB* gene cluster. We found the intact *raxX-raxSTAB* gene cluster exclusively in *Xanthomonas* spp., and ultimately detected it in more than 200 unique genome sequences (**File. S1**) among 413 accessed through the RefSeq database (O’Leary *et al.*, 2016).

*Xanthomonas* taxonomy has undergone several changes over the years (Vauterin *et al.*, 2000, Young, 2008) (see (Midha & Patil, 2014) for a representative example). At one point, many strains were denoted as pathovars of either *X. campestris* or *X. axonopodis*, but today over 20 species are distinguished, several with multiple pathovars (Rademaker *et al.*, 2005, Vauterin *et al.*, 1995). Because many of the genome sequences we examined are from closely-related strains, in some cases associated with different species designations, we constructed a whole-genome phylogenetic tree as described in Materials and Methods in order to organize these sequences by relatedness (**Fig. S1**). The topology of the resulting tree shares broad similarity with several other *Xanthomonas* phylogenetic trees in defining relationships between well-sampled species (Midha & Patil, 2014, Rademaker et al., 2005, Hauben *et al.*, 1997, Parkinson *et al.*, 2007, Parkinson *et al.*, 2009, Ferreira-Tonin *et al.*, 2012, Gardiner *et al.*, 2014, Triplett *et al.*, 2015, Young, 2008).

To examine *raxX-raxSTAB* gene cluster organization and inheritance more closely, we selected 15 genomes from strains that represent the phylogenetic range of *Xanthomonas* spp. (**Table 1** and **Fig. S1**). Where possible, we chose complete genome sequences that are accompanied by published descriptions. Throughout the analyses described below, species for which relatively large numbers of sequences are available also were monitored broadly for exceptional features. The close relative *Stenotrophomonas maltophilia*, which does not contain the *raxX-raxSTAB* gene cluster, serves as the outgroup (Moore *et al.*, 1997).

**Table 1.**
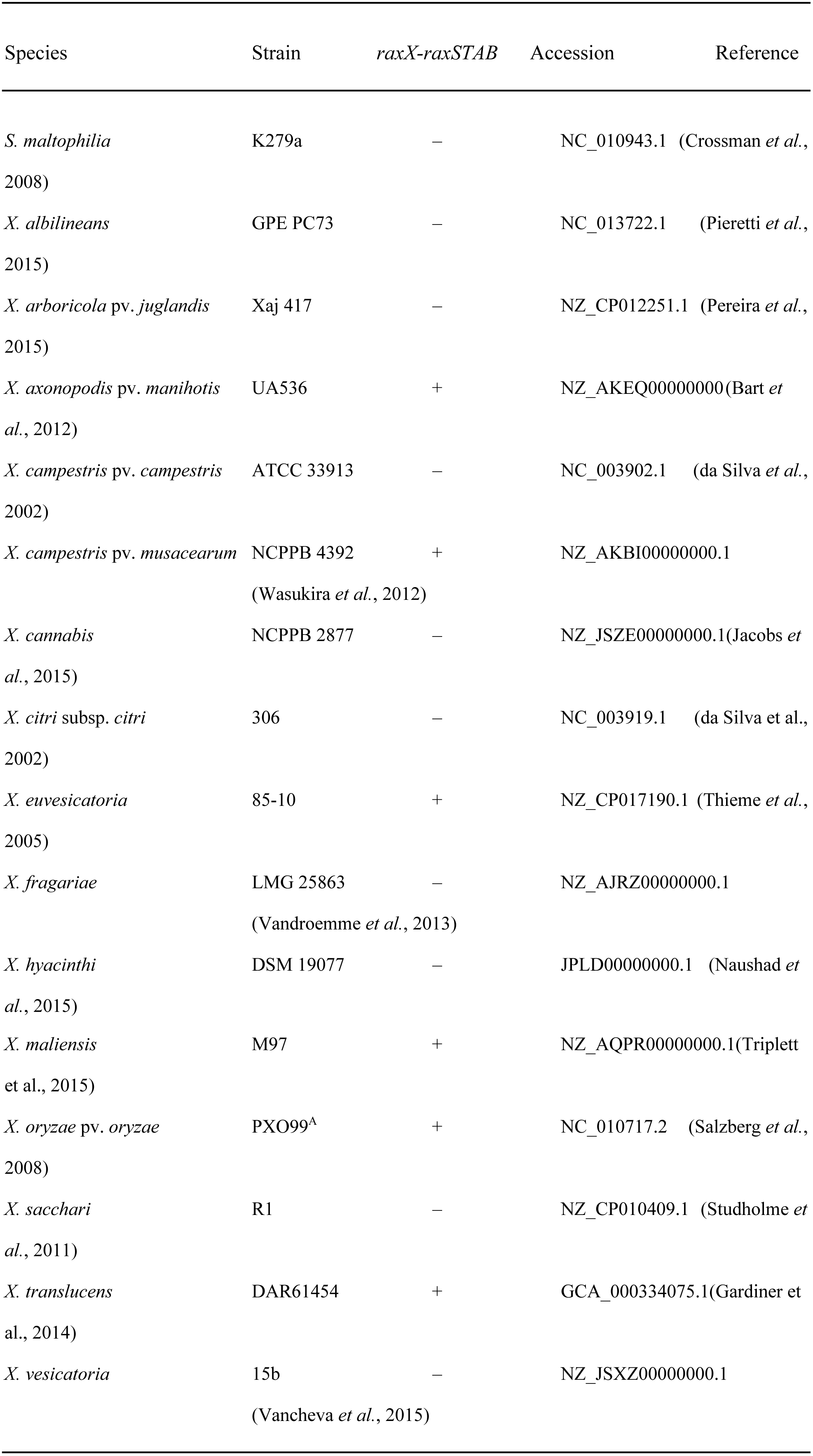
Reference strains for sequence comparisons.

To facilitate discussion, we represent phylogenetic relationships between these strains as a cladogram that emphasizes relative positions of the *raxX-raxSTAB* gene cluster-positive lineages (**Fig. 3**). Six distinct *Xanthomonas* lineages contain the *raxX-raxSTAB* gene cluster, one being *X. oryzae*. A second lineage includes related strains currently denoted as *X. vasicola* or *X. campestris* pv. *musacearum* (Aritua *et al.*, 2008); for concise presentation, we refer to these collectively as *X. vasicola*. The third lineage includes *X. euvesicatoria* and related species (Rademaker group 9.2; (Rademaker et al., 2005, Barak *et al.*, 2016). The fourth lineage includes strains denoted as *X. axonopodis*, such as pv. *manihotis* (Rademaker group 9.4; (Rademaker et al., 2005, Mhedbi-Hajri *et al.*, 2013). The fifth lineage includes *X. translucens* (Langlois *et al.*, 2017), within the distinct cluster of "early-branching" species whose divergence from the remainder apparently occurred relatively early during *Xanthomonas* speciation (Parkinson et al., 2007). The sixth lineage comprises *X. maliensis*, associated with but nonpathogenic on rice (Triplett et al., 2015). Phylogenetic analyses place this species between the "early-branching" species and the remainder (Triplett et al., 2015).

**Fig. 3.**
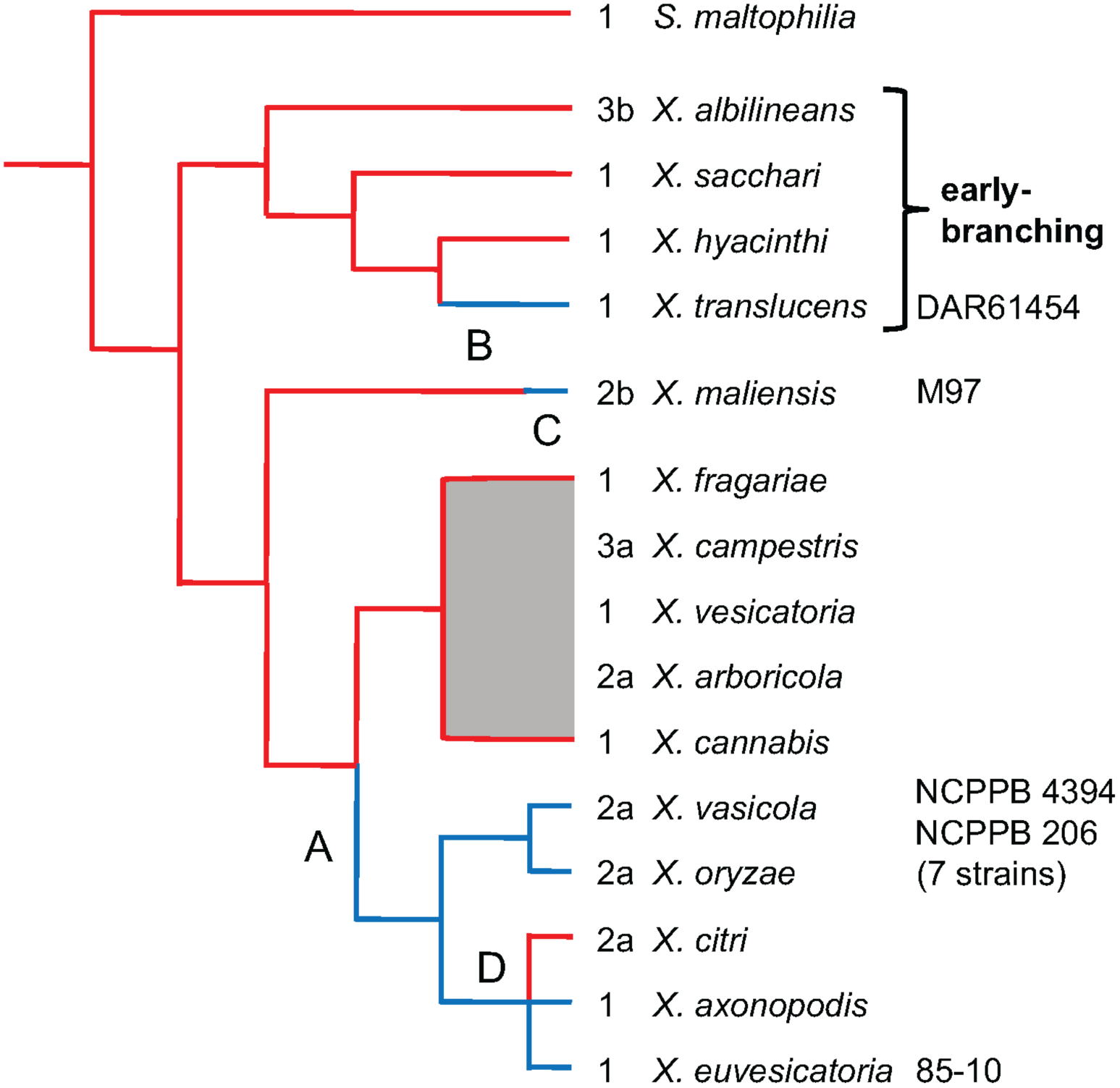
Model for raxX-raxSTAB inheritance during Xanthomonas speciation. The *Xanthomonas* spp. cladogram is based on published phylogenetic trees; see text for references. Red lines depict lineages for strains that lack the *raxX-raxSTAB* gene cluster, whereas blue lines depict those that carry the cluster. Numbers indicate *gcvP* length polymorphism in each species (see **Fig. S3**). Hypothetical events are: A, formation of the *raxX-raxSTAB* gene cluster; B, lateral transfer to *X. translucens*, relatively early during speciation (indicated by the long blue line); C, lateral transfer to *X. maliensis*, relatively late during speciation (indicated by the short blue line); D, loss from *X. citri*. Strain numbers denote sources of RaxX proteins chosen for functional tests, as described in the text.

Notably, the *raxX-raxSTAB* gene cluster is absent from the group of strains classified as *X. citri* pathovars (Rademaker group 9.5; (Rademaker et al., 2005, Bansal *et al.*, 2017). These strains (some of which are denoted as *X. axonopodis* or *X. campestris*) cluster phylogenetically among four of the *raxX-raxSTAB* gene cluster-positive groups: *X. oryzae*, *X. vasicola*, *X. euvesicatoria* and *X. axonopodis* pv. *manihotis* (Midha & Patil, 2014, Vauterin et al., 1995, Rademaker et al., 2005). The simplest explanation for this pattern is that the *raxX-raxSTAB* gene cluster was lost from an ancestor of the *X. citri* lineage (**Fig. 3**); other explanations are not excluded.

### Sequence conservation of the *raxX-raxSTAB* gene cluster suggests lateral transfer between *Xanthomonas* spp

Both the organization and size of the *raxX-raxSTAB* gene cluster are conserved across all six lineages. To assess inheritance patterns, we constructed a phylogenetic tree for the *raxX-raxSTAB* gene cluster (as the catenation of the four *rax* genes; **Fig. 4**) (Kuo & Ochman, 2009). The *rax* genes in *X. translucens*, in the early-branching group, cluster separately from their homologs in the other lineages. This finding is consistent with the hypothesis that *X. translucens* acquired the *raxX-raxSTAB* gene cluster relatively early during *Xanthomonas* speciation. For *X. maliensis*, the *raxX-raxSTAB* genes are most similar to those from *X. euvesicatoria* and the *X. axonopodis* pathovars *manihotis* and *phaseoli* (**Fig. 4**), even though the *X. maliensis* genome sequence itself is more distantly related (**Fig. 3**). This finding suggests that *X. maliensis* acquired the *raxX-raxSTAB* gene cluster relatively late during *Xanthomonas* speciation.

**Fig. 4.**
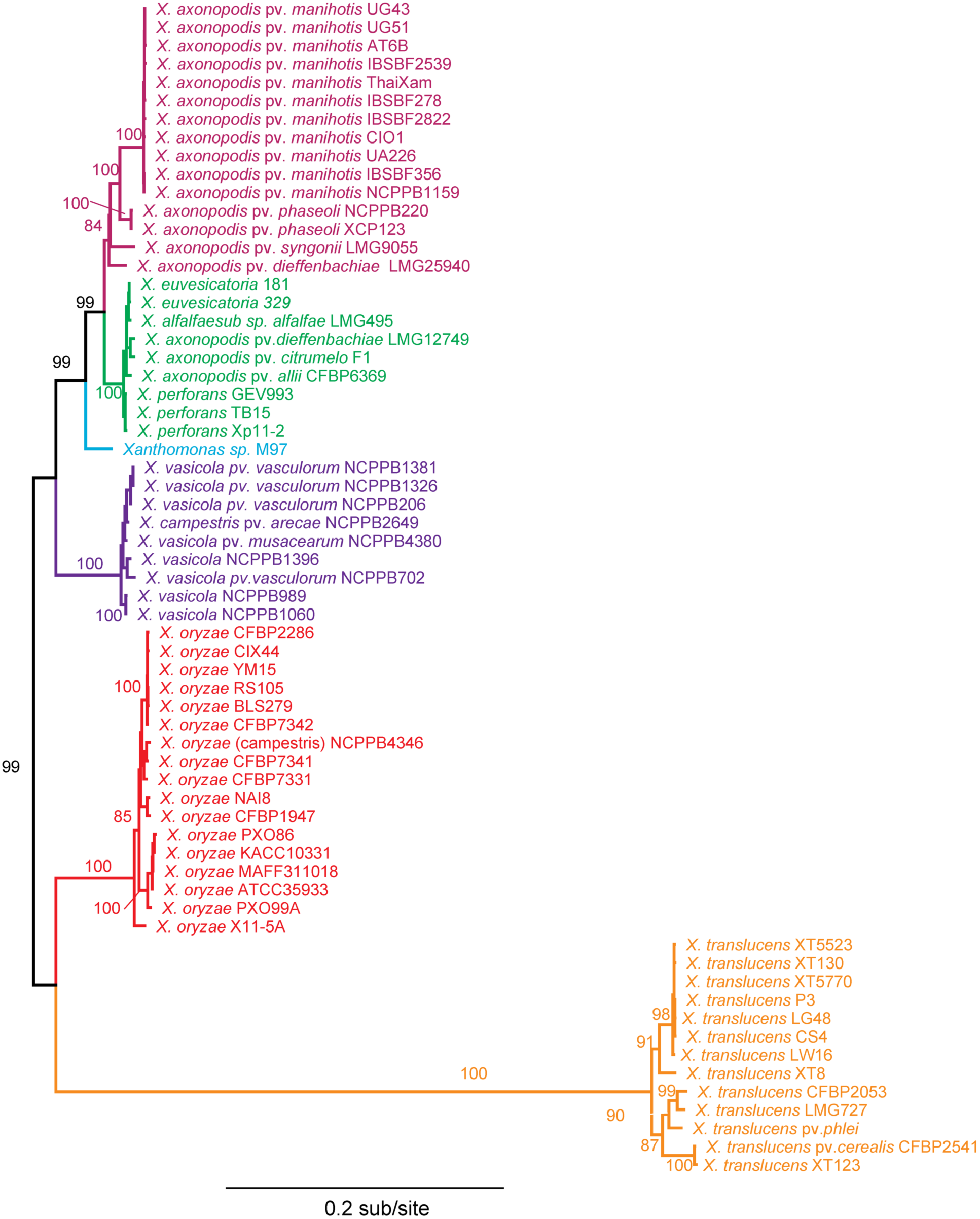
Phylogenetic tree for raxX-raxSTAB nucleotide sequences. The best scoring maximum likelihood tree for the catenated *raxA*, *raxB*, *raxX* and *raxST* coding sequences. Numbers shown on branches represent the proportion of branches supported by 10,000 bootstrap replicates (0-100). Bootstraps are not shown for branches with less than 50% support, nor for branches too short to easily distinguish. Species names are colored according to phylogenetic group.

### Boundaries flanking the *raxX-raxSTAB* gene cluster and adjacent genes suggest lateral transfer through general recombination

The *raxX-raxSTAB* gene cluster lies between two core (housekeeping) genes (**Fig. 1**). One, *gcvP*, encodes the pyridoxal-phosphate subunit of glycine dehydrogenase. An approximately 170 nt riboswitch (*gcvR* in **Fig. 1**) controls GcvP protein synthesis in response to glycine (Mandal *et al.*, 2004). The other, "*mfsX*", encodes a <u>**m**</u>ajor <u>**f**</u>acilitator <u>**s**</u>ubfamily (MFS) transporter related to Bcr and CflA efflux proteins (da Silva et al., 2004). Here, "*mfsX*" is only a provisional designation absent functional characterization.

Comparing the *gcvP - [raxX-raxSTAB]-* "*mfsX*" region from the reference genomes reveals sharp boundaries flanking the position of the *raxX-raxSTAB* gene cluster. On the left flank, substantial nucleotide identity spans the *gcvP* gene, the *gcvR* riboswitch, and a predicted *gcvR* promoter –10 element (Mitchell *et al.*, 2003) (**Fig. S2**). On the right flank, identity begins shortly after the "*mfsX*" initiation codon. Accordingly, upstream sequence elements for initiating "*mfsX*" gene transcription (Mitchell et al., 2003) and translation (Ma *et al.*, 2002) are conserved within, but not between, *raxX-raxSTAB* gene cluster-positive and -negative sequences (**Fig. S2**).

Between these boundaries in *raxX-raxSTAB* gene cluster-negative species, the compact (≤ 200 nt) *gcvP-*"*mfsX*" intergenic sequence is modestly conserved in most genomes (about 60-80% overall identity; **Fig. S2**). Much of this identity comes from the "*mfsX*" potential transcription and translation initiation sequences described above. The overall intergenic sequence is less conserved in the early-branching species (*X. albilineans*, *X. hyacinthi* and *X. sacchari*), displaying about 50-65% overall identity.

We hypothesize that *raxX-raxSTAB* gene cluster phylogenetic distribution results from general recombination between conserved genes flanking each side (e.g., in or beyond the *gcvP* and *"mfsX"* genes). Two observations are consistent with the hypothesis, First, we observed that the sequences flanking the *raxX-raxSTAB* gene cluster are different from the *gcvP-*"*mfsX*" intergenic sequence in *raxX-raxSTAB* gene cluster-negative strains (**Fig. S2**). This argues against models in which the *raxX-raxSTAB* gene cluster has integrated into the *gcvP-*"*mfsX*" intergenic sequence during lateral transfer events.

The second observation consistent with lateral transfer via general recombination is that *gcvP* length polymorphisms (**Fig 1** and **Fig. S3**) do not align with *Xanthomonas* phylogenetic relationships (**Fig. 3**). Inheritance patterns such as this often result from general recombination in the vicinity (Nelson *et al.*, 1997).

Notably, this *gcvP*-"*mfsX*" intergenic region conserved is also conserved in the *X. citr*i lineage (**Fig. S2**). If the *raxX-raxSTAB* gene cluster was lost during formation of this lineage (see above), then general recombination would replace the resident *raxX-raxSTAB* gene cluster with a donor conserved *gcvP*-"*mfsX*" region.

### *raxST* but not *raxX* homologs are present in genomes from diverse bacterial species

Our GenBank database searches identified *raxX* homologs and the *raxX-raxSTAB* gene cluster only in *Xanthomonas* spp. However, these searches did identify *raxST* homologs encoding proteins with about 40% identity to, and approximately the same length as, the *Xoo* RaxST protein. These sequences include the PAPS-binding motifs that define sulfotransferase activity (da Silva et al., 2004, Negishi *et al.*, 2001). Regardless of its current function, a *raxST* homolog potentially could evolve to encode tyrosylprotein sulfotransferase activity.

None of these *raxST* homologs is associated with a *raxX* homolog, and most also are not associated with *raxA* or *raxB* homologs. Presumably, the enzymes by these *raxST* homologs act on substrates other than RaxX. These *raxST* homologs support the hypothesis that the *raxSTAB* cluster arose from a new combination of pre-existing *raxST*, *raxA*, and *raxB* homologs. Proteolytic maturation and ATP-dependent peptide secretion systems are broadly distributed and so *raxA* and *raxB* homologs are plentiful in bacterial genomes (Holland et al., 2016).

These *raxST* homologs are in diverse genetic contexts in a range of bacterial phyla including Proteobacteria and Cyanobacteria (**Fig. S4**). Nevertheless, for most species represented by multiple genome sequences, the *raxST* homolog was detected in a minority of individuals, so it is not part of the core genome in these strains. Moreover, relationships between species in a *raxST* gene phylogenetic tree bear no resemblance to those in the overall tree of bacterial species. For example, in the *raxST* gene tree, sequences from Cyanobacteria are flanked on both sides by sequences from Proteobacteria (**Fig. S4**). Together, these findings provide evidence for lateral transfer of *raxST* homologs (Kuo & Ochman, 2009).

### RaxX protein sequence variants representing all six *raxX-raxSTA*B gene cluster-positive lineages

RaxX protein sequences from diverse *Xanthomonas* spp. assort into several sequence groups differentiated by polymorphisms within the predicted mature peptide sequence (**Fig. 2**) (Pruitt et al., 2017). Many of these groups are subdivided further according to polymorphisms in the predicted leader protein sequence (residues 1-39) or carboxyl-terminal region distal to residue Pro-52. Most leader polymorphisms lie between residues 2-24, and are unlikely to affect function of mature RaxX protein. Here we only consider polymorphisms in the predicted mature form.

To assess the function of RaxX variants, we focused on frequently observed variants in species represented by numerous genome sequences (**Fig. S1**). These include sequence groups A, B and D from *X. oryzae* pv. *oryzae* and *X. oryzae* pv. *oryzicola*, as well as sequence groups E, G and H, representing most genomes for the *X. euvesicatoria* and *X. vasicola* groups (**Fig. 2**). Finally, sequence group K is most numerous among *X. translucens* genomes. The comparison reference is the RaxX protein sequence from the Philippines *Xoo* strain PXO99A (sequence group A). Examples from lower frequency (mostly unique) sequence groups were analyzed by complementation, as described below.

### RaxX variants promote root growth but fail to activate the XA21-mediated immune response

We generated and purified tyrosine-sulfated full-length (unprocessed) RaxX peptides for these seven variants using an expanded genetic code approach (see methods) (**Fig. 2**), together representing all five pathogenic lineages that contain the *raxX-raxSTAB* gene cluster. The positive control is RaxX21-sY, a synthetic 21 residue tyrosine-sulfated peptide with strong activity (Pruitt et al., 2015). These peptides were used in two separate assays for function. First, we performed root growth experiments with an *Arabidopsis thaliana tpst-1* mutant lacking tyrosylprotein sulfotransferase, which is required for all known sulfated tyrosine peptide hormones including PSY (Matsubayashi, 2014). This eliminates endogenous PSY activity so that effects of added peptides are more easily observed (Pruitt et al., 2017, Matsubayashi, 2014). Root lengths for seedlings grown without added peptide averaged 23.5 mm, whereas root lengths for seedlings grown with 100 nM peptide were at least twice as long (**Fig. 5A** and **Fig. 5B**). This observation is consistent with the hypothesis that these peptides mimic PSY1 peptide hormone activity. Note that three of these variants (groups D, E and G) were examined previously (Pruitt et al., 2017) and are included here to facilitate direct comparisons as well as to monitor consistency of results.

**Fig. 5.**
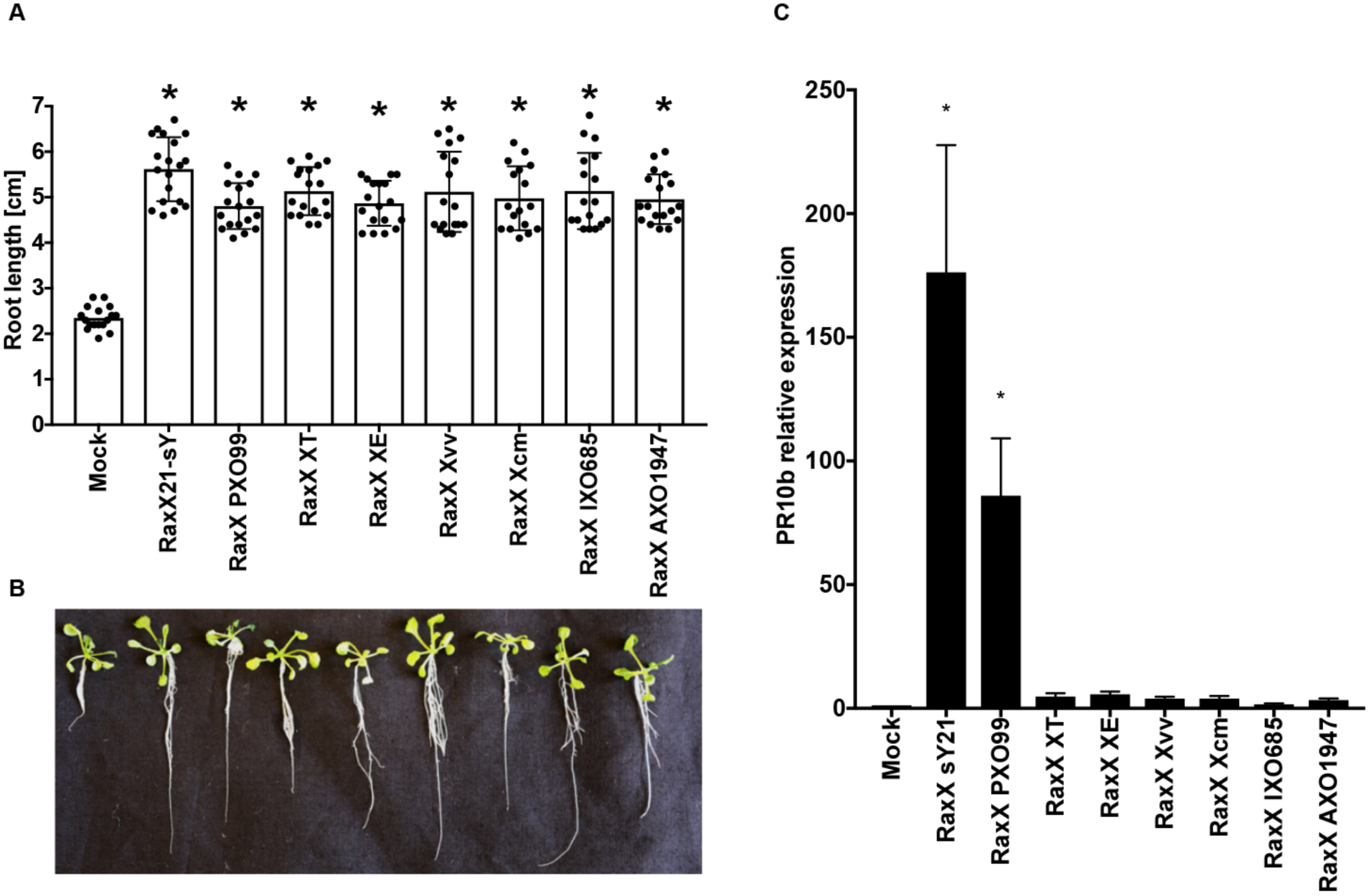
RaxX variant peptides promote root growth. (A) Stimulation of *Arabidopsis* root growth. Fourteen-day-old *tpst-1* seedlings were grown on ½ MS vertical plates with or without 100 nM of the indicated full-length peptides. Bars indicate the average seedling root length measured after 14 d (n>10). Error bars show the standard deviation. The “*” indicates a statistically significant difference from Mock using Dunnett’s test (p<0.05). Peptide RaxX sY21 is a 21 residue sulfated peptide with potent RaxX activity (Pruitt et al., 2015). Strain abbreviations are *Xvv, X. vasicola* pv. *vasculorum*; *Xt*, *X. translucens*; *Xe*, *X. euvesicatoria*; *Xcm*, *X. campestris* pv. *musacearum*; PXO99^A^, IXO685, AXO1947, strains of *X. oryzae* pv. *oryzae*. (B) *Arabidopsis seedlings* from a representative experiment. (C) Activation of rice *PR10b* gene expression. Purified peptide (500 nM) was used to treat detached leaves as described in Materials and Methods. Expression levels of the *PR10b* gene (normalized to actin gene expression) were determined after 12 h. Data are the mean values from four biological replicates. Error bars show the standard deviation. The “*” indicates a statistically significant difference from Mock using Dunnett’s test (*p*<0.05).

In the second assay, we tested each RaxX peptide for direct activation of XA21-mediated immunity by assaying induction of the *PR10b* marker gene as a readout for immune activation (Thomas *et al.*, 2016, Pruitt et al., 2015). In contrast to results with the root growth assay, here only the group A RaxX protein (from *Xoo* strain PXO99A) was able to induce XA21-mediated *PR10b* marker gene expression (**Fig. 5C**).

In a separate test for activation of XA21-mediated immunity, we used a □*raxX* deletion mutant of *Xoo* strain PXO99^A^ as a host for genetic complementation. We tested each of the *raxX* alleles shown in **Fig. 2**, which includes examples from lower frequency (mostly unique) sequence groups. We introduced each *raxX* allele into the □*raxX* test strain, and monitored disease progression in leaves of whole plants. Only the group A *raxX* allele (from *Xoo* strain PXO99A) was able to complement the *Xoo* PXO99^A^ □*raxX* strain to activate XA21-mediated immunity (**Fig. 6**). Expression of each *raxX* allele was confirmed by qPCR (**Fig. S5**).

**Fig. 6.**
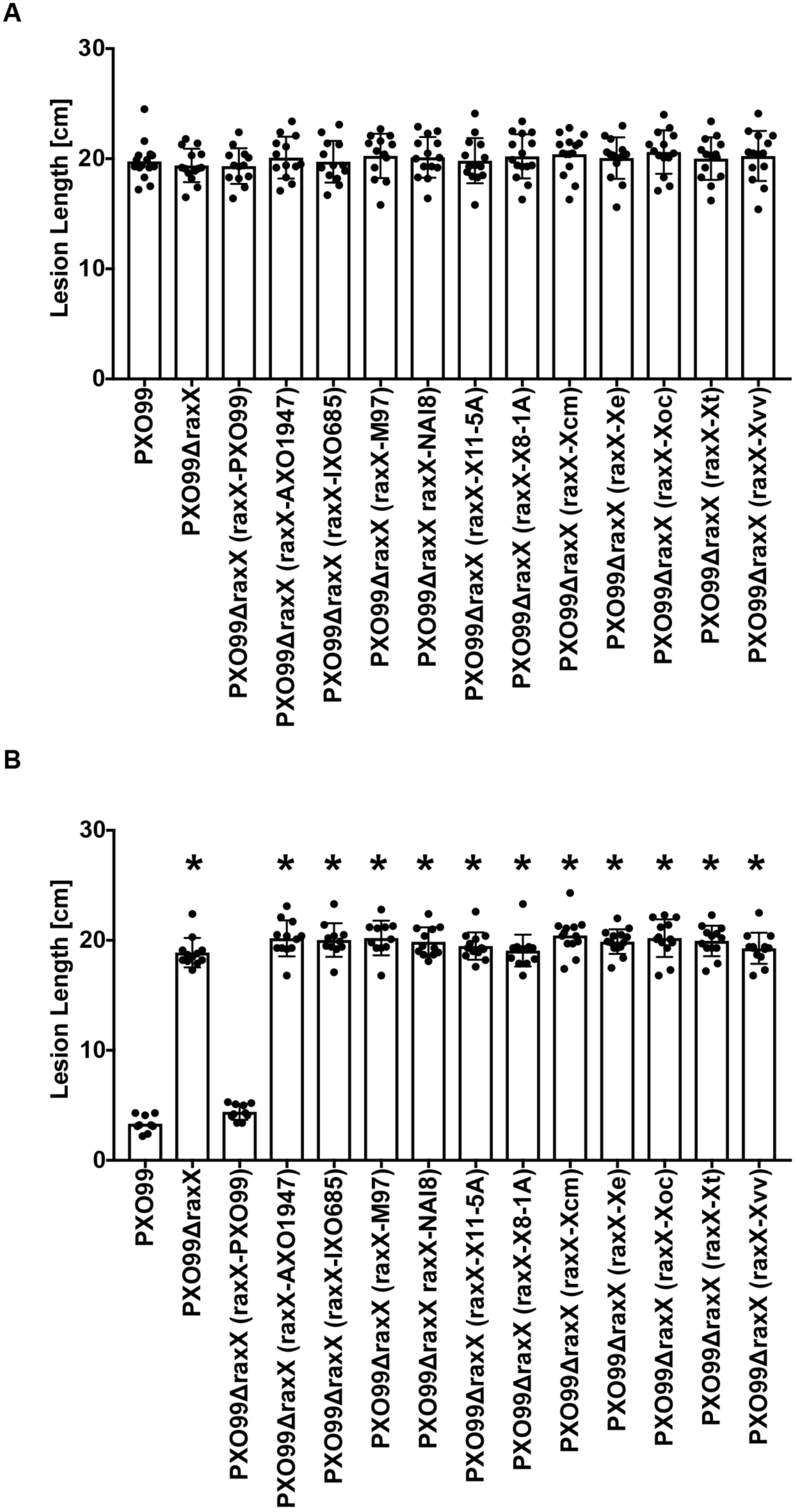
RaxX variants fail to activate XA21-mediated immunity. Different *raxX* genes were cloned into vector pVSP6 (see Materials and Methods) to test for complementation of the *Xoo* strain PXO99^A^ ∆*raxX* strain. Leaf tips of rice varieties TP309 (panel A) or XA21-expressing TP309 (panel B) were inoculated by clipping with scissors dipped in bacterial suspensions (approximate cell density of 8 × 10^8^ cells mL^-1^). Lesion lengths were measured 14 days after inoculation. Data are the mean values from measurements of 10-20 leaves. Error bars show the standard error of the mean, and “*” indicates a statistically significant difference from *Xoo* strain PXO99^A^ according to Dunnett’s multiple comparison procedure (*p*<0.05). Values in panel A are insignificantly different. Strain abbreviations are Xvv, *X. vasicola* pv. *vasculorum*; Xt, *X. translucens*; Xoc, *X. oryzae* pv. *oryzicola*; Xe, *X. euvesicatoria*; Xcm, *X. campestris* pv. *musacearum*; X8-1A, X11-5A, strains of *X. oryzae*; M97, *X. maliensis* M97; PXO99^A^, IXO685, AXO1947, strains of *X. oryzae* pv. *oryzae*.

Together, these results provide direct evidence that activation of XA21-mediated immunity is restricted to RaxX proteins from sequence group A, found in most strains of *Xoo*. None of the other *X. oryzae* RaxX variants tested (including RaxX from *X. oryzae* pv. *oryzicola*) was able to activate XA21-mediated immunity. The observation that all RaxX proteins tested stimulated *Arabidopsis* root growth suggests that the RaxX PSY peptide mimicry function is not restricted to rice.

### African *Xoo* strain AXO1947 RaxX and RaxST natural variants both lead to evasion of the XA21 immune receptor

The *raxX* alleles from *Xoo* strains IXO685 and AXO1947 failed to complement the □*raxX* mutant of *Xoo* strain PXO99^A^ for XA21 immune activation (**Fig. 6**). In addition to its variant *raxX* allele (**Fig. 2**), we noted that *Xoo* strain AXO1947 (Huguet-Tapia *et al.*, 2016) carries seven missense polymorphisms in the *raxST* gene (**Fig. S6**) not present in other *Xoo* strains such as IXO685. To determine if the variant *raxST* allele from strain AXO1947 encodes a functional protein, we performed additional complementation tests.

We found that the *raxX* allele from strain PXO99^A^ conferred the XA21 immune activation phenotype upon strain IXO685 but not upon strain AXO1947 (**Fig. 7B**). This result suggests that the *raxX* variant allele is not the only factor that prevents strain AXO1947 from activating the XA21 immune response. Consistent with this hypothesis, the *raxST* allele from strain PXO99^A^ failed to confer the XA21 immune activation phenotype upon strain AXO1947 (**Fig. 7D**). In contrast, addition of both the *raxX* and *raxST* alleles from strain PXO99^A^ was sufficient to confer the XA21 immune activation phenotype upon strain AXO1947 (**Fig. 7F**).

**Fig. 7.**
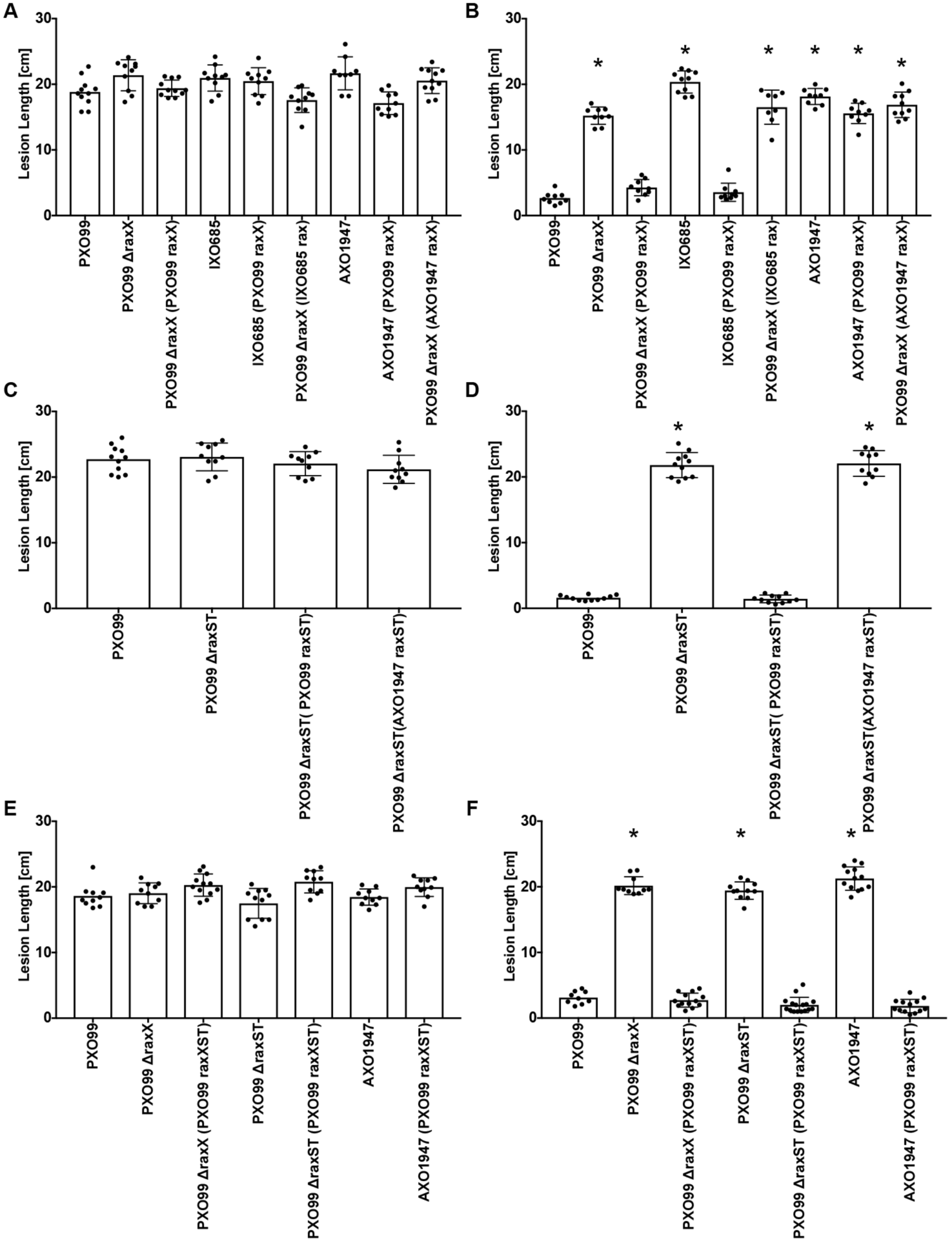
The *raxX* and *raxST* genes are dysfunctional in *Xoo* strain AXO1947. Different combinations of the *raxX* and *raxST* genes were cloned into vector pVSP61 (see Materials and Methods) to test for complementation. Leaf tips of rice varieties TP309 (panels A, C and E) or XA21-expressing TP309 (panels B, D and F) were inoculated by clipping with scissors dipped in bacterial suspensions (approximate cell density of 8 × 10^8^ cells mL^-1^). Lesion measurements were taken 14 days after inoculation. Data are the mean values from measurements of 10-20 leaves. Error bars show the standard error of the mean, and “*” indicates a statistically significant difference from *Xoo* strain PXO99^A^ according to Dunnett’s multiple comparison procedure (*p*<0.05). Values in panels A, C and E are insignificantly different. Panels A and B show complementation results for the *raxX* gene, panels C and D show results for the *raxST* gene, and panels E and F show results for the combination of both the *raxX* and *raxST* genes. Specific combinations of genes and complementation hosts are described in the figure labels.

Taken together, these results suggest that *Xoo* strain AXO1947 has mutant versions of both genes, *raxST* and *raxX*. Analysis by qRT-PCR confirms that these genes were expressed in the complemented strains (**Fig. S7**).

### RaxST variants from *Xoo* strain AXO1947

To determine which of the RaxST missense polymorphisms is responsible for the apparent reduction in enzyme activity, we used site-specific mutagenesis to introduce each individually into the *raxST* gene from strain PXO99^A^. Genes encoding two of these [His-50 to Asp (H50D) and Arg-129 to Leu (R129L)] were unable to complement the □*raxST* mutant of *Xoo* strain PXO99^A^ for XA21 immune activation (**Fig. 8**), indicating that both His-50 and Arg-129 are necessary for RaxST activity.

**Fig. 8.**
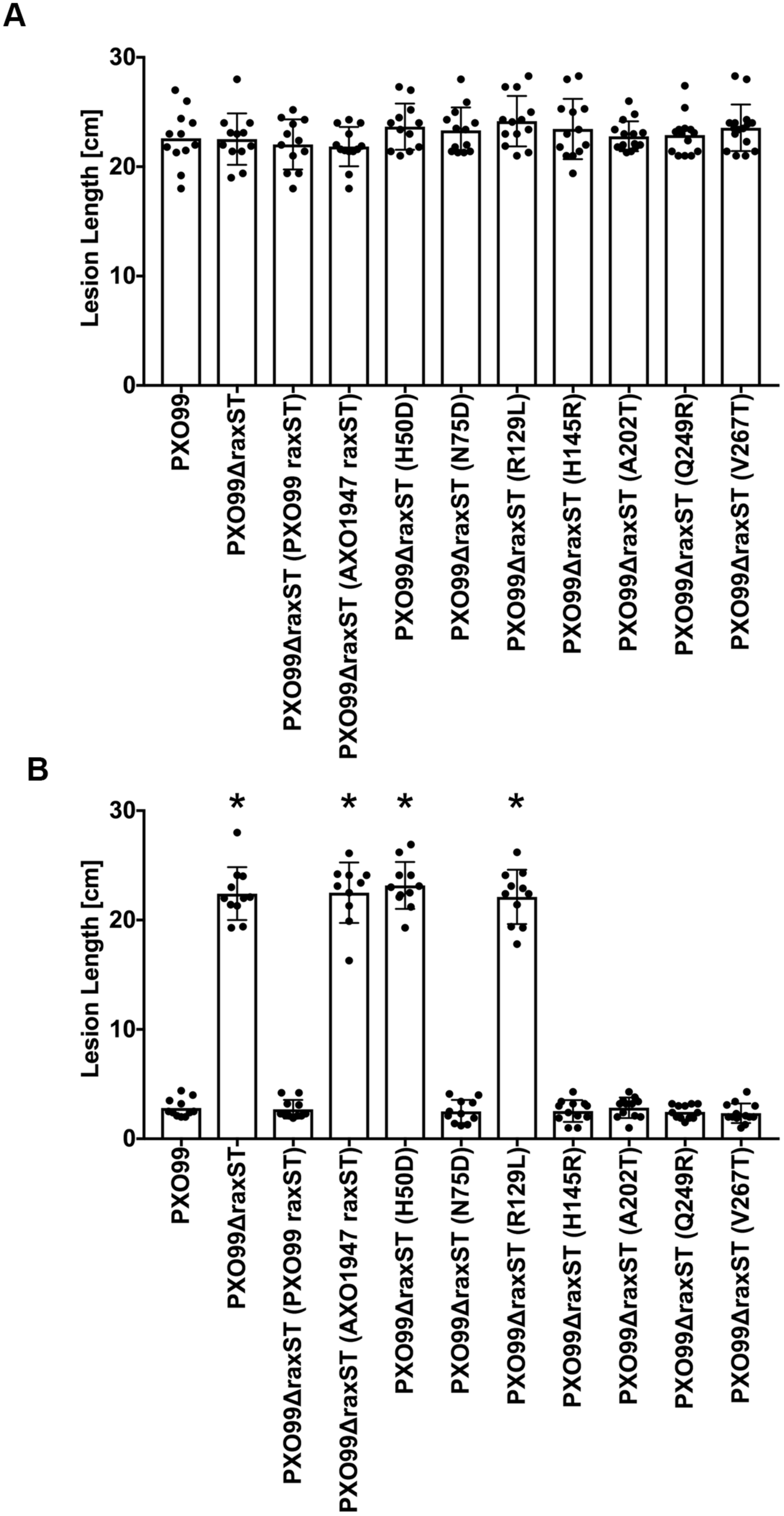
Two missense substitutions inactivate RaxST in *Xoo* strain AXO1947. Each of the seven *raxST* missense polymorphisms from *Xoo* strain AXO1947 was introduced singly into the wild-type *raxST* gene from *Xoo* strain PXO99^A^ (see Materials and Methods). These mutant alleles then were tested for complementation of the *Xoo* strain PXO99^A^ ∆*raxST* strain. Leaf tips of rice varieties TP309 (panel A) or XA21-expressing TP309 (panel B) were inoculated by clipping with scissors dipped in bacterial suspensions (approximate cell density of 8 × 10^8^ cells mL^-1^). Lesion measurements were taken 14 days after inoculation. Data are the mean values from measurements of 10-20 leaves. Error bars show the standard error of the mean, and “*” indicates a statistically significant difference from *Xoo* strain PXO99^A^ according to Dunnett’s multiple comparison procedure (*p*<0.05).

Little is known about RaxST structure and function. Diverse sulfotransferases share limited sequence similarity, mostly comprising two relatively short sequence motifs involved in PAPS binding (Negishi et al., 2001). These motifs are conserved in the *Xoo* RaxST sequence (da Silva et al., 2004). Research with diverse sulfotransferases has identified three essential residues: a positively-charged residue (corresponding to Arg-11 in RaxST) in one PAPS binding motif, an invariant Ser (corresponding to Ser-118 in RaxST) in the other, and a catalytic base (His or Glu) located between the two PAPS binding motifs (Negishi et al., 2001).

We generated a RaxST molecular model with the program iTasser (Yang & Zhang, 2015) using the crystal structure of human tyrosylprotein sulfotransferase-2 (TPST2) as a template (PDB: 3AP1). The sequence alignment is shown in **Fig. S8**. TPST2 is a functional dimer (Teramoto *et al.*, 2013), which is replicated in the RaxST structural model (**Fig. S9)**. The two essential residues identified from *Xoo* strain AXO1947, His-50 and Arg-129, display surface exposed side chains in close proximity to the corresponding position for the bound substrate peptide co-crystalized with TPST2. These residues are distal to the catalytic site. Therefore, we hypothesize that these RaxST residues are involved in RaxX peptide binding.

## DISCUSSION

We previously hypothesized that RaxX mimics the actions of PSY hormones, and that the XA21 receptor evolved specifically to recognize RaxX from *Xoo* (Pruitt et al., 2015, Pruitt et al., 2017). This prediction is supported here by our finding that all the RaxX variants tested stimulate root growth (**Fig. 5A** and **Fig. 5B**) (Pruitt et al., 2017) but fail to activate the XA21-mediated immune response (**Fig. 5C** and **Fig. 6**). Thus, RaxX sequence determinants are more stringent for XA21-mediated immunity activation than for root growth stimulation. In this discussion, we consider two questions: (1) What are potential selective pressures acting on RaxX that affect sequence variation; and (2) How was the *raxX-raxSTAB* gene cluster inherited in *Xanthomonas* spp.?

### Opposing selection pressures drive RaxX natural variation

Maintenance of the *raxX-raxSTAB* gene cluster (**Fig. 3**) suggests that RaxX provides fitness benefits to diverse *Xanthomonas* spp., presumably during their interactions with hosts that collectively encompass a range of monocot and dicot species. This hypothesis is supported by *in vivo* data showing that *Xoo* strains lacking the *raxX* or *raxST* genes are compromised for virulence (Pruitt et al., 2015, Pruitt et al., 2017). On the other hand, rice-restricted XA21-mediated immunity would select specifically against RaxX maintenance by *Xoo*. Analysis of *raxX-raxSTAB* gene cluster sequence polymorphisms suggests that both types of selection occur.

The *Xa21* gene has been introgressed into commercial rice varieties (Khush *et al.*, 1990, Midha *et al.*, 2017). Widespread planting of *Xa21* rice presumably increases selection for *Xoo* variants that evade XA21-mediated immunity. All RaxX missense variants examined mimicked PSY hormone activity (**Fig. 5A** and **Fig. 5B**) (Pruitt et al., 2017), suggesting that this property confers a selective advantage. Consistent with this, we did not observe any *raxX* frameshift or nonsense alterations. Instead, RaxX variant sequences contain a restricted set of missense substitutions, consistent with the hypothesis that selection acts to retain PSY-like function (**Fig. 2**; see reference (Pruitt et al., 2017)).

Among all RaxX variants tested, only that from *Xoo* strain PXO99A (which represents the large majority of *Xoo raxX* alleles) activated the XA21-mediated immune response (**Fig. 5C** and **Fig. 6**). This result demonstrates that recognition of RaxX by XA21 is strictly limited to *Xoo*, and confirms and extends a prior conclusion from our laboratory, that residues Pro-44 and Pro-48 both are required for *Xoo* RaxX recognition by XA21 (Pruitt et al., 2015).

Thus, it appears that some *Xoo* strains that evade activation of XA21-mediated immunity arise from a restricted set of *raxX* missense substitution alleles encoding variants that retain PSY-like function. This observation suggests that it may be possible to engineer novel XA21 variants that recognize these variant RaxX proteins. If so, it may then be possible to engineer broad-spectrum resistance against *Xoo* (and other *raxX-raxSTAB* gene cluster-positive *Xanthomonas* spp.) by expressing multiple XA21 proteins that collectively recognize multiple RaxX variants.

We also have identified *raxST* and/or *raxA* gene loss of function alterations in *Xoo* field isolates (**Fig. 7**; reference (da Silva et al., 2004), which presumably cannot express the PSY mimicry phenotype of RaxX). Such loss of function alterations could temper the effectiveness of production strategies that rely on engineered *Xa21* alleles.

### *raxX-raxSTAB* gene cluster origin

The *raxAB* genes are homologous to those encoding proteolytic maturation and ATP-dependent peptide secretion complexes (da Silva et al., 2004, Lin *et al.*, 2015), related to type 1 secretion systems but specialized for secreting small peptides such as bacteriocins and peptide pheromones (Holland et al., 2016). Frequently, the gene encoding the secreted substrate is adjacent to genes encoding components of the secretion complex (Dirix *et al.*, 2004). We hypothesize that the intact *raxX-raxSTAB* gene cluster originated in an ancestor to the lineage containing *X. oryzae*, *X. euvesicatoria*, and related species, with subsequent gains or loss through lateral transfer (**Fig. 2**). Relatively few events appear to have been necessary to form the *raxX-raxSTAB* gene cluster. The *raxX* gene might have evolved from the gene for the secreted peptide substrate of the RaxAB ancestor. The complete cluster would result from incorporation of the ancestral *raxST* gene, homologs of which are distributed broadly (**Fig. S4**).

### Role for the *raxX-raxSTAB* gene cluster in *Xanthomonas* biology

The *raxX-raxSTAB* gene cluster does not exhibit features, such as a gene for a site-specific recombinase, characteristic of self-mobile genomic islands (Hacker *et al.*, 1997). Instead, evidence suggests that *raxX-STAB* gene cluster lateral transfer occurred through general recombination between genes flanking each side of the *raxX-STAB* gene cluster (**Fig. 1** and **Fig. S2**). In bacteria, gene acquisition through lateral transfer contributes to emergence of new pathovars (see reference (Ogura *et al.*, 2009) for one example). Conceivably, lateral acquisition of the *raxX-raxSTAB* gene cluster might allow a particular strain to infect a previously inaccessible host.

*Xanthomonas* pathovar phenotypes (Jacques *et al.*, 2016) are not predicted by the presence or absence of the *raxX-raxSTAB* gene cluster. For example, some *raxX-raxSTAB* gene cluster-positive species can infect only monocots (e.g., *X. oryzae*, *X. translucens*) or only dicots (e.g., *X. euvesicatoria*), just as some *raxX-raxSTAB* gene cluster-negative species also can infect only monocots (e.g., *X. arboricola*, *X. hyacinthi*) or only dicots (e.g., *X. campestris* pv. *campestris*; *X. citri*). Similarly, some *raxX-raxSTAB* gene cluster-positive species are specific for vascular tissue (e.g., *Xoo*; *X. vasicola*) or for non-vascular tissue (e.g., *X. oryzae* pv. *oryzicola*; *X. euvesicatoria*), just as some *raxX-raxSTAB* gene cluster-negative species also are specific for vascular tissue (e.g., *X. hortorum*; *X. albilineans*) or for non-vascular tissue (e.g., *X.citri*; *X. arboricola*). Thus, selective function(s) for the *raxX-raxSTAB* gene cluster in *Xanthomonas* spp. remain to be determined.

### Experimental Procedures

#### Survey of the RaxX, RaxST and the *raxX-STAB* genomic region in publicly available databases

We used the 5kb long *Xoo* PXO99^A^ *raxX-raxSTAB* genomic region, including 600 bp upstream of *raxST* and 70 bp downstream of *raxB*, as query to search the following NCBI databases with blastn and megablast using e-value cut-off of 1e-3; nr/nt, htgs, refseq_genomic_representative_genomes, refseq_genomic, and gss. To identify RaxX homologs we used the protein sequence of RaxX from *Xoo* PXO99^A^ as query to search the same databases using tblastn with a PAM30 scoring matrix to account for the short sequence length of RaxX. In case of *raxST* from *Xoo* PXO99^A^ we used the genomic coding sequence to search the same databases using the same cut-offs. In addition, we used the RaxST protein sequence to search the following database using blastp with an e-value cut-off of 1e-3 and a BLOSUM62 scoring matrix; nr, refseq_protein, env_nr. The databases were last accessed 2016/01/06 for the initial manuscript submission and 2018/06/25 during preparation of the resubmission. Searches were restricted to bacteria (taxid: 2) in case of refseq_genomic_representative_genomes. The observations of specificity of *raxX* and the intact *raxX-raxSTAB* gene cluster to the genus *Xanthomonas* was consistent across all queries.

#### Whole genome based phylogenetic tree for *Xanthomonas* spp

All available *Xanthomonas* genomes were downloaded from the NCBI ftp server on January 29, 2016 (413 genome accessions). The genome fasta files were used to build a local blast database using BLASTv2.27+ (Camacho *et al.*, 2009). For all genes in and surrounding the *raxSTAB* cluster blastn (evalue cutoff of 1e-3) was used to identify homologs in the local blast database. Due to the small size of RaxX, tblastn was required to identify homologs (evalue cutoff of 1e-3). Fasta files for each blast hit were generated using a custom python script (available upon request). Alignments of all genes were performed with Muscle v3.5 (Edgar, 2004) implemented in the desktop tool Geneious v9.1.8 (Kearse *et al.*, 2012). Alignment ends were trimmed so that each sequence was equal in length and in the first coding frame. Maximum likelihood trees were built with RaxML v8.2.4 (Stamatakis, 2014) with the following settings: (-m GTRGAMMA F -f a -x 3298589 -N 10000 -p 23). Trees shown in all figures are the highest scoring ML tree and numbers shown on branches are the resampled bootstrap values from 1000 replicates. Trees were drawn in FigTree v1.4.0 (http://tree.bio.ed.ac.uk/software/figtree/).

Whole genome phylogenies were generated using the entire genome assembly with the program Andi v0.10 (Haubold *et al.*, 2015, Klotzl & Haubold, 2016). These distance matrices were plotted as neighbor-joining tree using Phylip v3.695 (Felsenstein, 1981). Numbers on the branches represent the proportion (0-100) that the branch appeared in the “bootstrapped” distance matrices using Andi.

#### Sequence analyses

Nucleotide and deduced amino acid sequences were edited and analyzed with the programs EditSeq^TM^ (version 14.1.0), MegAlign^TM^ (version 14.1.0) and SeqBuilder^TM^ (version 14.1.0), DNASTAR, Madison, WI. The Integrated Microbial Genomes interface (Chen *et al.*, 2017) was used to compare genome segments from different species.

#### Bacterial growth

*Xanthomonas* strains were cultured at 28°C. Solid medium was peptone sucrose agar (PSA; pH 7.0), which contains (per liter) peptone (10 g), sucrose (10 g), sodium glutamate (1 g) and agar (15 g). Liquid cultures were aerated at 230 rpm in YEB medium (pH 7.3), which contains (per liter) yeast extract (5 g), tryptone (10 g), NaCl (5 g), sucrose (5 g), and MgSO_4_ (0.5 g). Antibiotics were kanamycin, carbenicillin, spectinomycin (all at 50 □g/ml), and cephalexin (20 □g/ml).

#### Rice growth and inoculation

*Oryza sativa* ssp. *japonica* rice varieties were TP309 and XA21-TP309, which is a 106-17-derived transgenic line of TP309 carrying the *Xa21* gene expressed from its native promoter (Song et al., 1995). TP309 rice does not contain the *Xa21* gene. Seeds were germinated in distilled water at 28°C for one week and then transplanted into sandy soil (80% sand, 20% peat; Redi-Gro) in 5.5-inch square pots with two seedlings per pot. Plants were grown in tubs in a greenhouse, and were top watered daily with fertilizer water [N, 58 ppm (parts per million); P, 15 ppm; K, 55 ppm; Ca, 20 ppm; Mg, 13 ppm; S, 49 ppm; Fe, 1 ppm; Cu, 0.06 ppm; Mn, 0.4 ppm; Mo, 0.02 ppm; Zn, 0.1 ppm; B, 0.4 ppm] for four weeks, followed by water for two weeks. Six weeks after planting, rice pots were transferred to a growth chamber with the following day/night settings: 28°C/24°C, 80%/85% humidity, and 14/10-hour lighting. Plants were inoculated 2 to 3 days after transfer using the scissors clipping method (Song et al., 1995). Bacteria for inoculation were taken from PSA plates and resuspended in water at a density of approximately 8 × 10^8^ CFU/ml. Water-soaked lesions were measured 14 days after inoculation.

#### Complementation tests

The *Xoo* strain PXO99^A^ marker-free deletions □*raxX* and □*raxST* were described previously (Pruitt et al., 2015). The *raxX* and *raxST* genes from different *Xanthomonas* spp. were cloned into plasmid vector pVS61 and electrotransformed into the appropriate recipient strains as described previously (Pruitt et al., 2015). Site-specific mutational alterations were introduced by PCR using the In-Fusion HD cloning system (Takara).

#### RaxX peptide stimulation of *PR10b* gene expression

Full-length sulfated RaxX proteins were purified from an *E. coli* strain with an expanded genetic code that directs incorporation of sulfotyrosine at the appropriate position (Schwessinger *et al.*, 2016). The resulting MBP-3C–RaxX-His fusion proteins were incubated with 3C protease followed by anion exchange chromatography in order to remove the amino-terminal maltose binding protein tag, as described previously (Schwessinger et al., 2016). The control peptide, sulfated RaxX21-sY, has been described (Pruitt et al., 2015).

Rice plants were grown in a hydroponic system in growth chambers at 24° or 28°C with a 14-hour/10-hour light-dark cycle at 80% humidity. Seedlings were grown in A-OK Starter Plugs (Grodan) and watered with Hoagland’s solution twice a week. Peptide influence on *PR10b* marker gene expression was measured as described previously (Pruitt et al., 2015). Briefly, leaves of 4-week-old hydroponically grown rice plants were cut into 2-cm-long strips and incubated for at least 12 hours in ddH_2_O to reduce residual wound signals. Leaf strips were treated with the indicated peptides and then snap-frozen in liquid nitrogen before processing. Quantitative reverse transcription polymerase chain reaction (qRT-PCR) was done as described previously (Pruitt et al., 2015). Gene expression was normalized to the actin gene expression level and to the respective mock-treated control at 0 or 9 hour.

DNA primers for qRT-PCR were: ampC-F, GACTCGTAATGCCTACGACC; ampC-R, AATTGCTCGTAGAAGCTGCC; qraxST-F, CTTCCAACGTGCAGATCGAC; qraxST-R, TATCGACGATCCAACCAAC; qRaxX-F, AAAATCGCCCGCCAAGGGT; qRaxX-R, TCAATGGTGCCCGGGGTTG; PR10b-F, TGTGGAAGGTCTGCTTGGAC; PR10b-R, CCTTTAGCACGTGAGTTGCG

## Acknowledgements

Supported by NIH GM59962 and GM122968 to Pamela C. Ronald. We are grateful to anonymous reviewer 2 for homology modeling. We thank Simon Williams from Research School of Biology at Australia National University, Canberra and Michael Steinwand from the Ronald laboratory for their assistance with RaxST homology modeling.

## Supporting Information

**Fig. S1.**
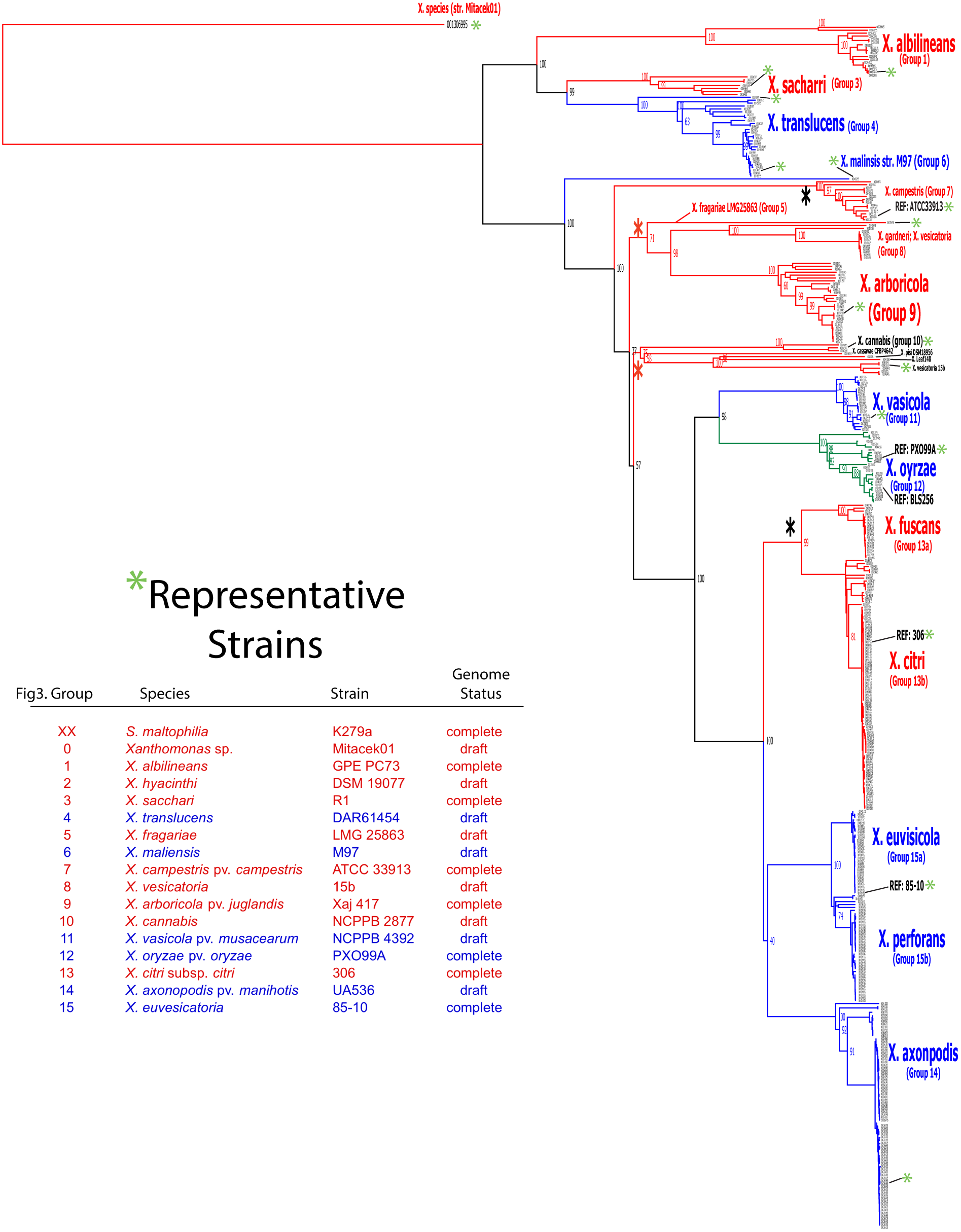
Whole genome-based *Xanthomonas* phylogenetic tree. This tree was constructed by analysis of whole genome sequences as described in Materials and Methods. Blue indicates genomes that contain the *raxX-raxSTAB* gene cluster; red indicates genomes that do not. Group numbers are arbitrary.

**Fig. S2.**
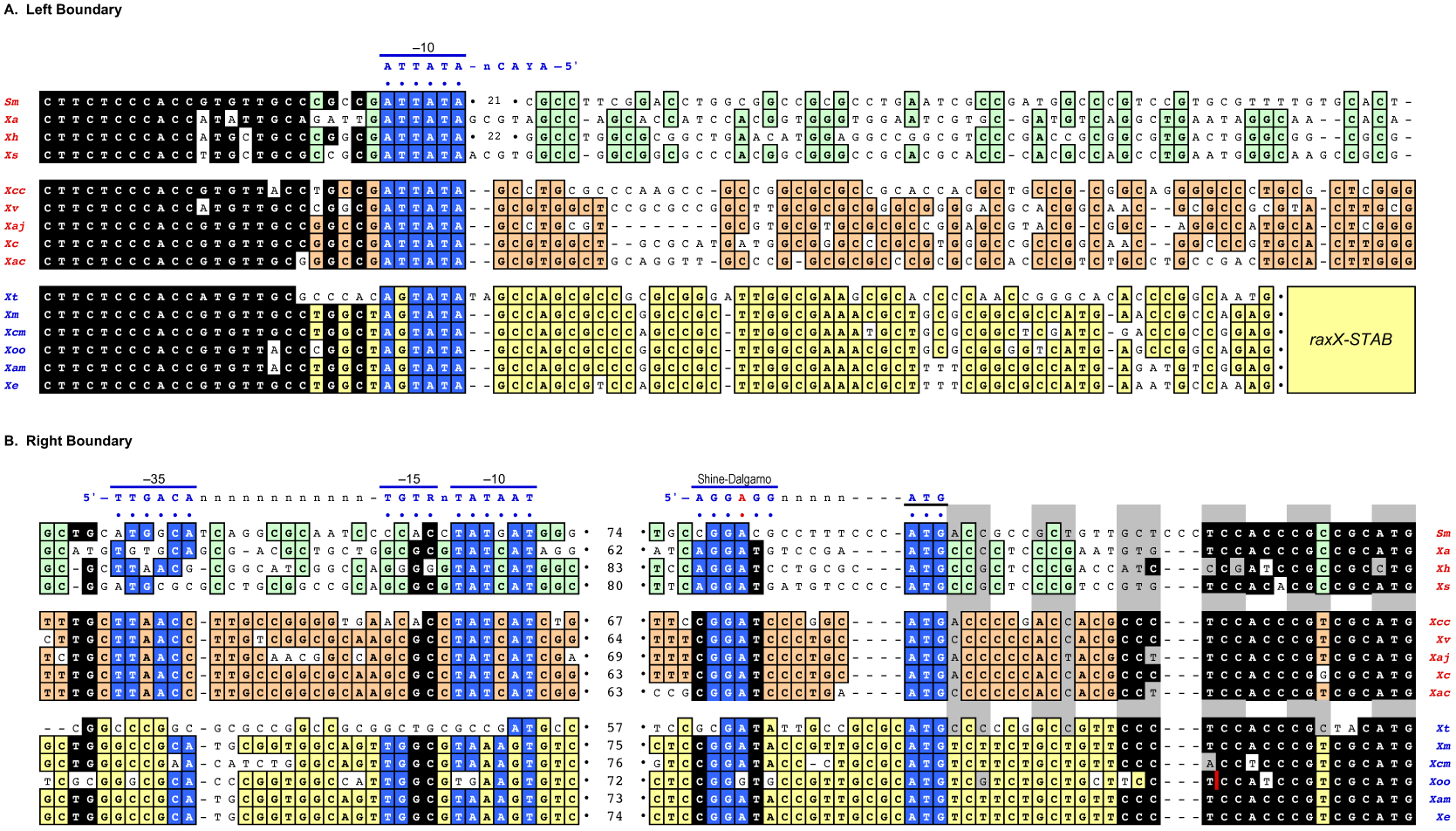
Sequences flanking the *raxX-raxSTAB* gene cluster. Sequences are from the reference strains described in **Table 1**. Sequences conserved within a group but different from other groups are colored green ("early-branching" species), brown (*raxX-raxSTAB* cluster-negative strains), or yellow (*raxX-raxSTAB* cluster-positive strains). For presentation, the sequence is divided into Left and Right boundaries. The green and brown sequences are contiguous, whereas the yellow sequences are interrupted by the ca. 5 kb *raxX-raxSTAB* gene cluster, depicted as a yellow rectangle. For presentation, approximately 60-80 nt with relatively low similarity were removed from sequence shown in the Right boundary panel. These conceptual deletions are denoted by the number of nt removed in each case. Black sequences are conserved in all lineages, and include both coding regions as well as matches to transcription and translation initiation consensus sequences, which are described in the text. An “*mfsX*” +1 frameshift in *Xoo* sequences is indicated by the vertical red line. Abbreviations are in red for *raxX-raxSTAB* cluster-negative strains and blue for *raxX-raxSTAB* cluster-positive strains: *S. maltophilia, Sm; X. albilineans, Xa; X. arboricola pv. juglandis, Xaj; X. axonopodis pv. manihotis, Xam; X. campestris pv. campestris, Xcc; X. campestris pv. musacearum, Xcm; X. cannabis, Xc; X. citri subsp. citri, Xac; X. euvesicatoria, Xe; X. fragariae, Xf; X. hyacinthi, Xh; X. maliensis, Xm; X. oryzae pv. oryzae,Xoo; X. sacchari,; Xs X. translucens, Xt; X. vesicatoria, Xv.*

**Fig. S3.**
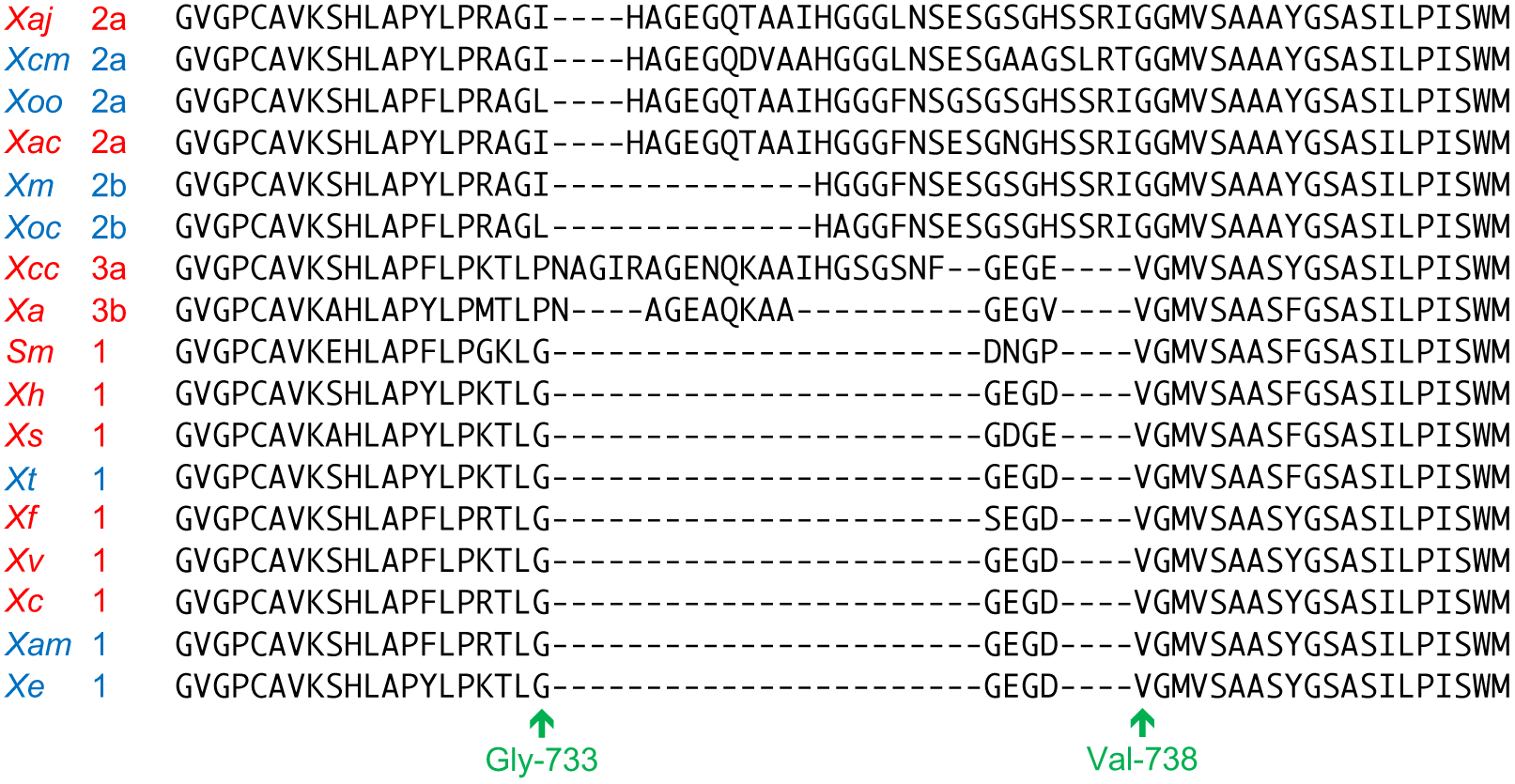
GcvP length polymorphisms in different *Xanthomonas* lineages. The relevant portion of the GcvP amino acid sequence is shown for each of the reference strains. Species in red lack the *raxX-raxSTAB* gene cluster, whereas those in blue carry the cluster. Numbers denote different allelic types for reference to **Fig. 3**. The positions of residues Gly-733 and Val-738 (numbering for allelic type 1) are indicated. Abbreviations: *S. maltophilia, Sm; X. albilineans, Xa; X. arboricola pv. juglandis, Xaj; X. axonopodis pv. manihotis, Xam; X. campestris pv. campestris, Xcc; X. campestris pv. musacearum, Xcm; X. cannabis, Xc; X. citri subsp. citri, Xac; X. euvesicatoria, Xe; X. fragariae, Xf; X. hyacinthi, Xh; X. maliensis, Xm; X. oryzae pv. oryzae,Xoo; X. sacchari,; Xs X. translucens, Xt; X. vesicatoria, Xv.*

**Fig. S4.**
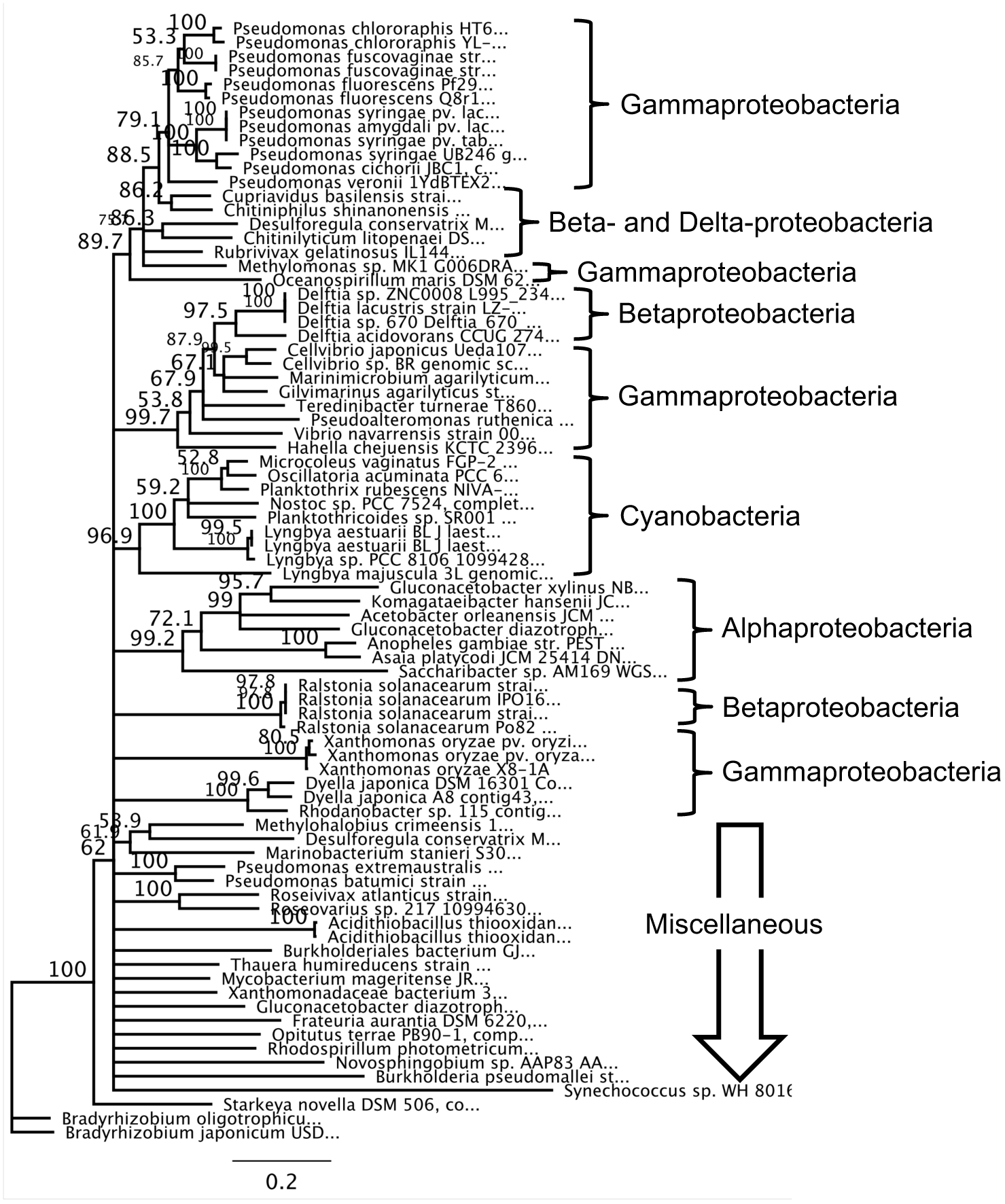
Phylogenetic tree for *raxST* homologs. Distribution of *raxST* homologs across bacterial genera, including the major groups of proteobacteria as well as cyanobacteria. The tree shown was constructed by neighbor-joining with 1000 bootstrap replicates; branches with < 50% bootstrap support are not drawn. The *raxST* sequence from *Xoo* strain PXO99A was used as query for tBLASTn.

**Fig. S5.**
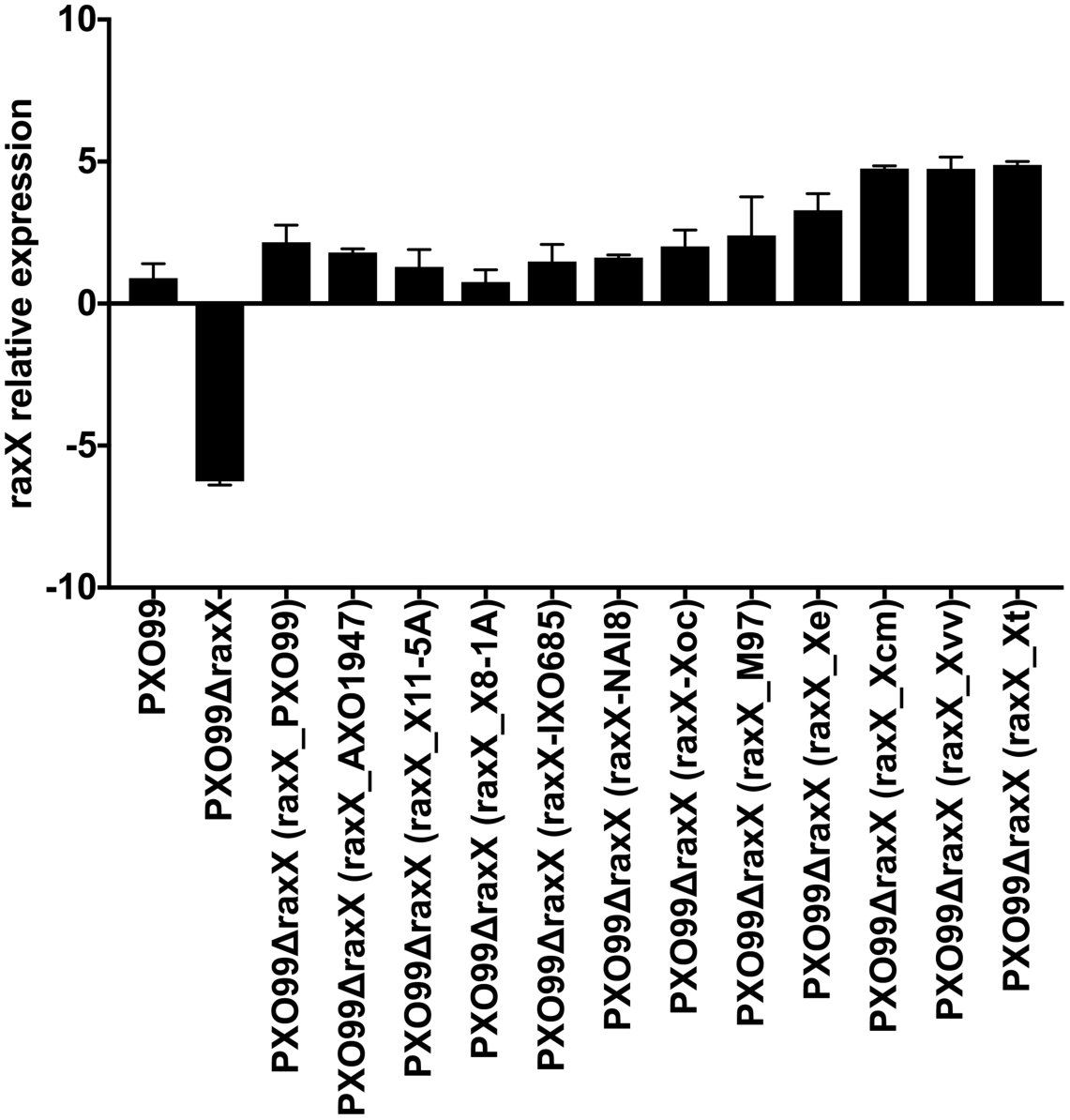
*raxX* expression in *Xoo* PXO99^A^ complemented strains. Data show that *raxX* gene expression in the complemented strains with different *raxX* alleles with its promoter region on plasmids. The expression is shown as the logarithm of raw data using qRT-PCR. Gene expression was normalized to the chromosomal gene PXO_01660 (annotated as an *ampC* gene homolog encoding ß-lactamase). Data are the mean values from two biological replicates. Error bars show the standard deviation.

**Fig. S6.**
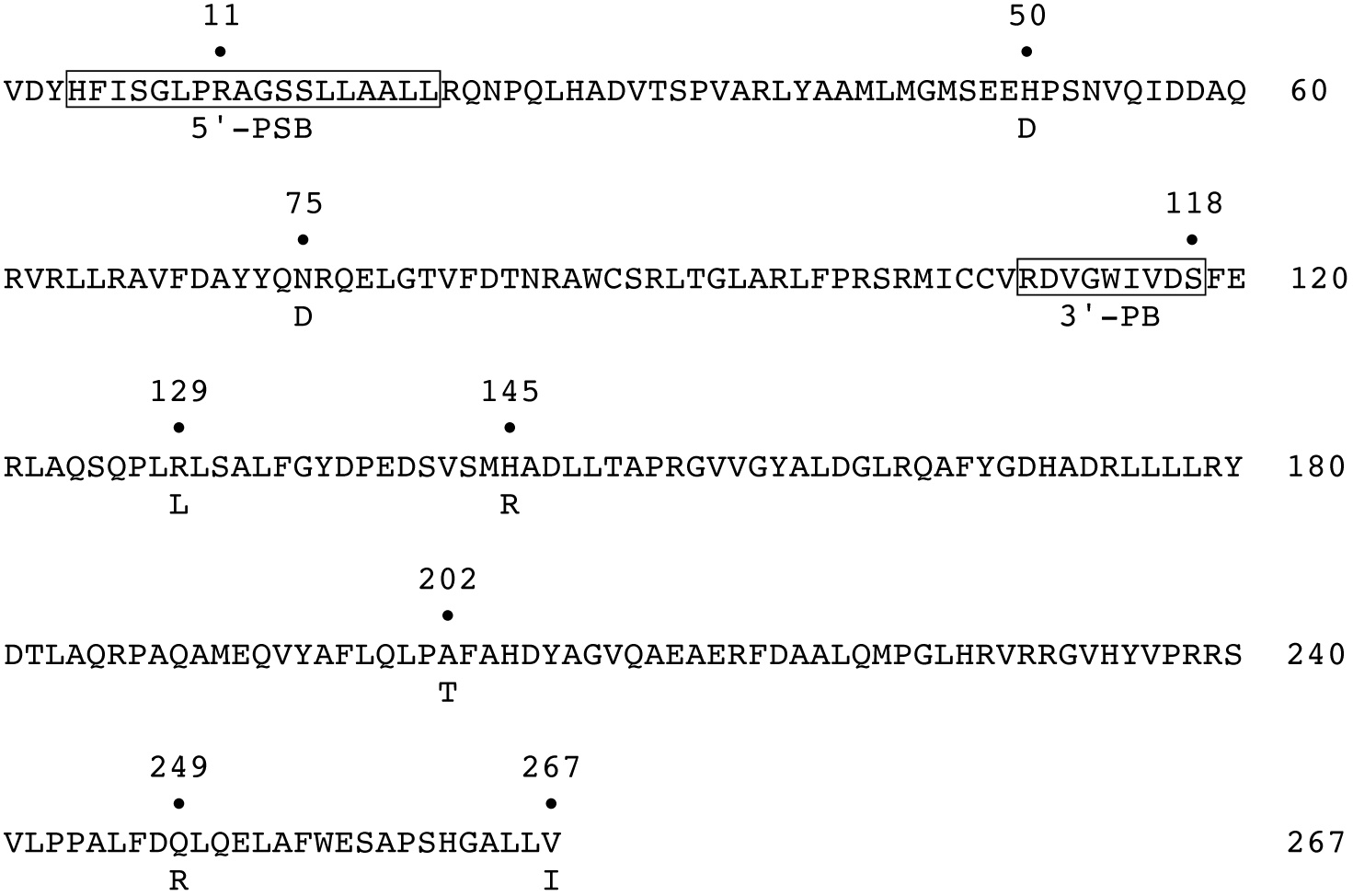
RaxST sequence polymorphisms in *Xoo* strain AXO1947. The RaxST sequence from *Xoo* strain PXO99^A^ is shown. The seven missense substitutions in the sequence from *Xoo* strain AXO1947 (Huguet-Tapia et al., 2016) are indicated. The boundaries of the PAPS binding motifs (5’-PSB and 3’-PB; reference (Negishi et al., 2001), are enclosed in boxes. These motifs encompass the catalytic residues Arg-11 and Ser-118.

**Fig. S7.**
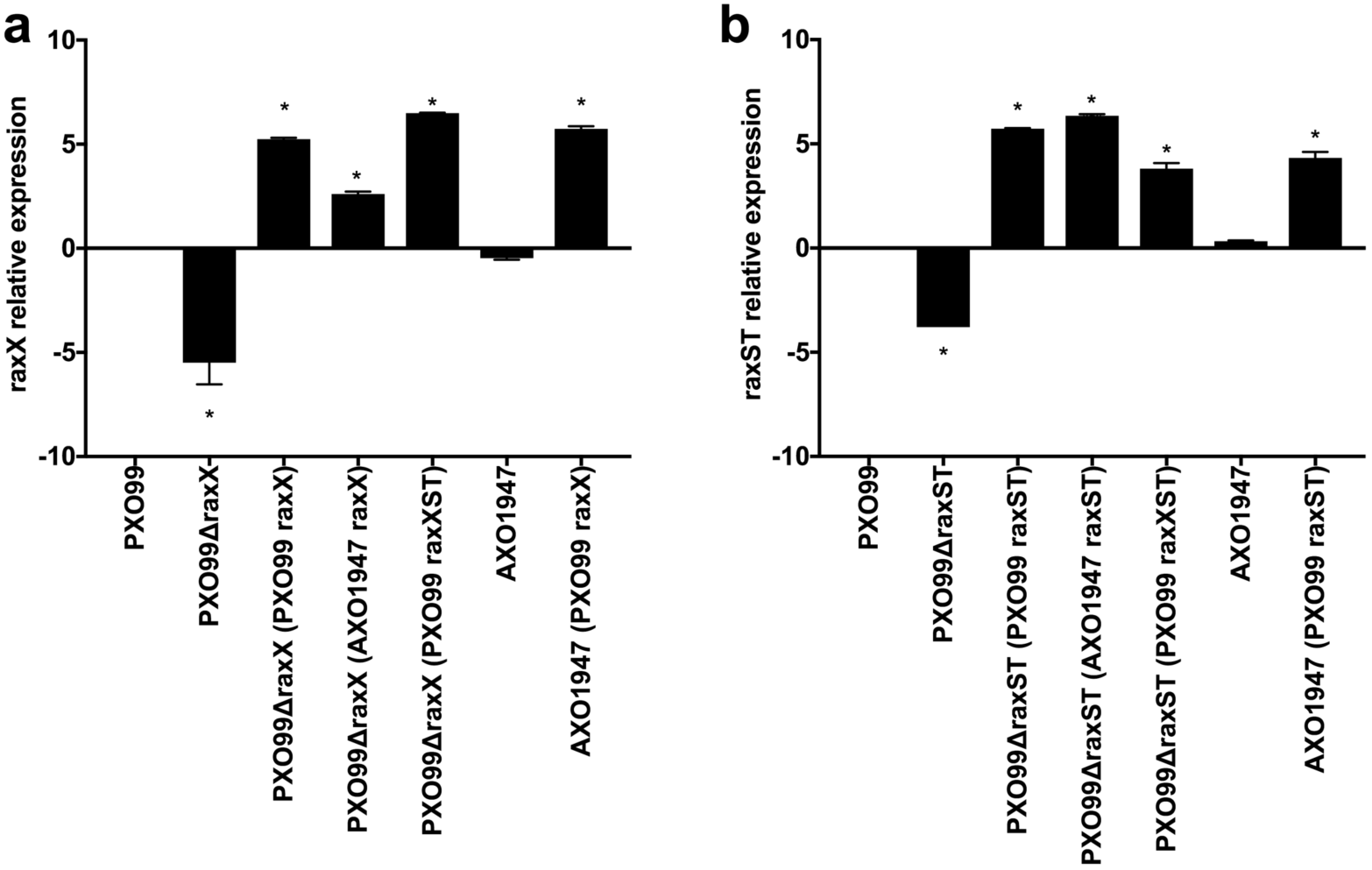
*raxX* and *raxST* expression in *Xoo* PXO99^A^ complemented strains. Data show *raxX* and *raxST* gene expression in the complemented strains (with *raxX* and *raxST* on plasmids) relative to expression in *Xoo* strain PXO899^A^ (with *raxX* and *raxST* on the chromosome). Expression was determined by qRT-PCR (see Materials and Methods), and is shown as the logarithm of the fold change. Gene expression was normalized to the chromosomal gene PXO_01660 (annotated as an *ampC* gene homolog encoding □-lactamase). Data are the mean values from two biological replicates. Error bars show the standard deviation.

**Fig. S8.**
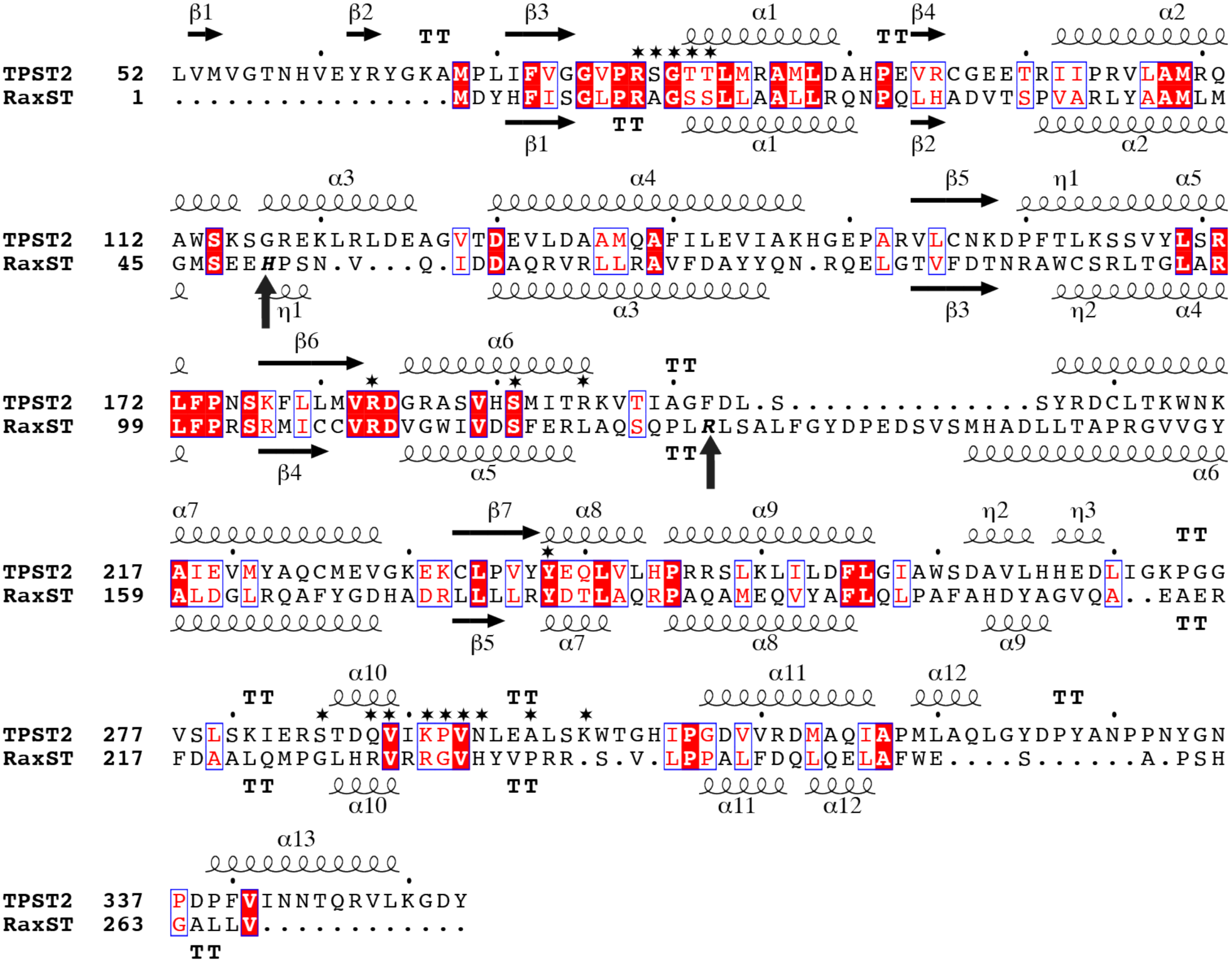
RaxST structural alignment. Sequence alignment of the human TPST2 and *Xoo* RaxST sequences formatted with ESPript 3.0 (Robert & Gouet, 2014). Secondary structure elements derived from the respective structural models are shown. Stars show TPST2 residues involved in PAPS binding, and arrows show RaxST missense substitutions.

**Fig. S9.**
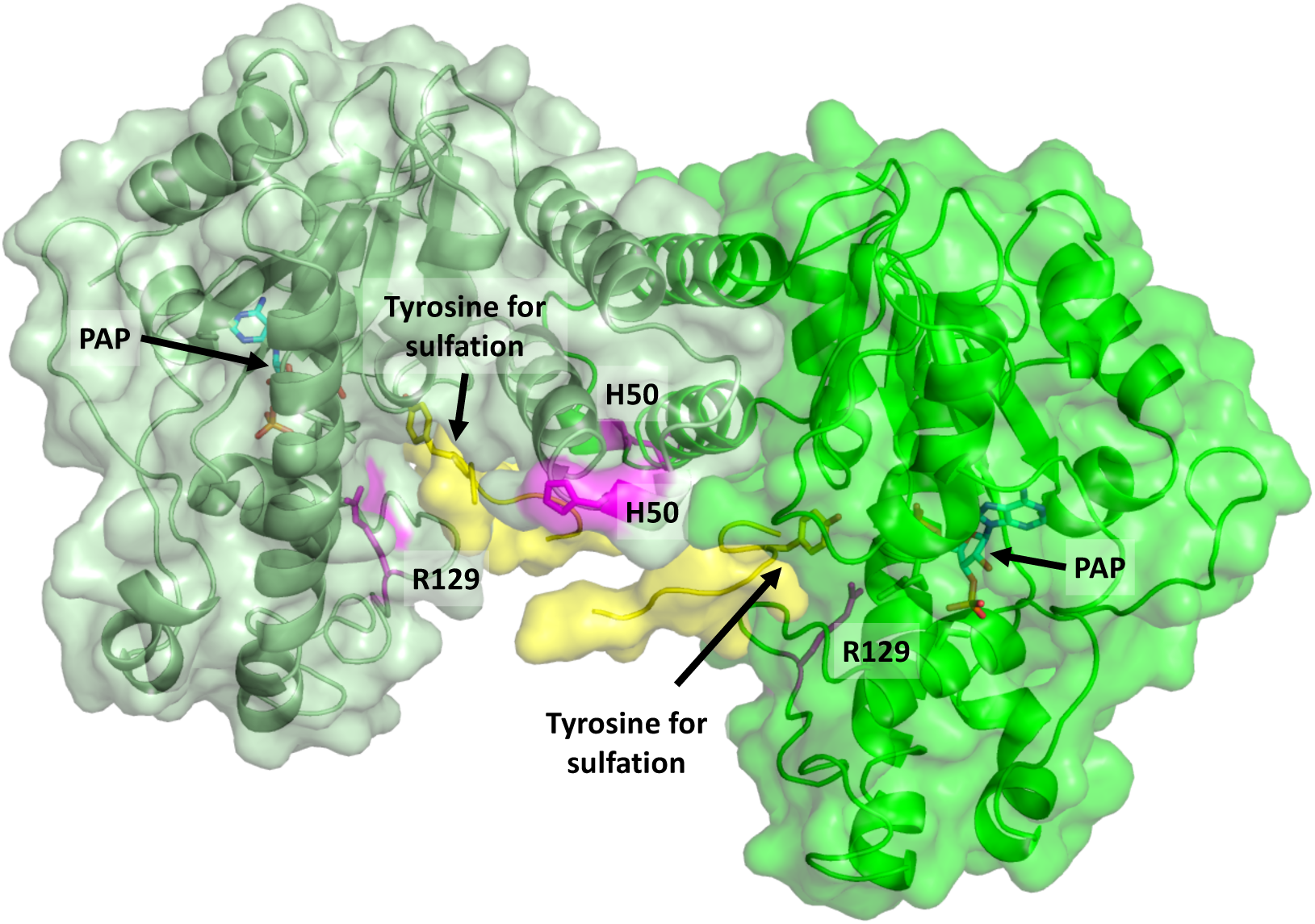
Model for RaxST structure. Predicted RaxST structure shown in cartoon and surface representation, based on the dimeric structure of TPST2. The two RaxST monomers are colored in dark and light green. The 3’-phosphoadenosine-5’-phosphate (PAP) and C4 substrate peptide that where co-crystallized with TPST2 are superimposed on the RaxST model. PAP is represented as labelled and the substrate peptide is shown in yellow-cartoon with the acceptor tyrosine represented as labelled. Residues His-50 and Arg-129 are colored in magenta and highlighted.

**File. S1. *Xanthomonas* strains analyzed for whole-genome phylogeny.**

Excel file (.XLS format).

